# Hematopoietic proliferation is orchestrated by the sequential and lineage-specific activation of Cyclin D, Cyclin E and CDKN sub-modules within the G1/S network

**DOI:** 10.1101/2025.05.15.654268

**Authors:** Andrea Hanel, Aya Abdelsalam, Sylvain Tollis

## Abstract

Commitment to the cell division cycle constrains other fate choices at the single cell level. Hence the molecular network controlling the G1/S transition must be coordinated with developmental phases. Healthy hematopoiesis relies on shifts in cell cycle dynamics that balance proliferation and differentiation of hematopoietic stem cells (HSCs) and lineage progenitors, offering an ideal model system to study the coordination of cell division with development. The G1/S network orchestrates this process by regulating the activites of the master G1/S transcription factors (E2F), and cyclin dependent kinases whose activities drive the progression to S phase. By using single cell transcriptomics profiles of human bone marrow cells and mathematical modeling, we demonstrate that variations in the expression of cyclin D- and cyclin E-centered sub-modules of the G1/S network carve out distinct trajectories from G1 to S, explaining the distinct proliferation properties of hematopoietic cell types evolving in the same microenvironment, and biasing cell fate decisions towards certain lineages. We map 68 hematopoietic cell types to specific model parameters, and identify their individual route through G1/S, using our model. This improved mechanistic understanding of the G1/S transition across cell types could enhance the design of more nuanced pharmacological strategies, enabling more personalized treatment recommendations for current cell cycle-targeted cancer therapies.

## Introduction

In all eukaryotic species, cells commit to division by activating a large transcriptional program at the G1/S transition of the cell division cycle (reviewed in (Bertoli, Skotheim, et de Bruin 2013)). The molecular network that controls this transcriptional burst is highly conserved – from plants (Vercruysse et al. 2020), to yeasts (Hendler et al. 2018), and mammals (Bertoli, Skotheim, et de Bruin 2013). The regulation of the cell cycle, and the G1/S commitment point in particular, is among the most extensively studied pathways in cell biology, owing to its central relevance in e.g. cancer and regenerative medicine. Over the past 70 years, intensive research has identified the key proteins and their interactions that together form the G1/S network. Although the network architecture is well described, it is way less understood from a functional perspective: in particular, it is not known which upstream signals trigger the activation of the G1/S transcriptional programs; nor we can explain why cells that cohabit the same microenvironment, and are therefore exposed to the same external stimuli, exhibit markedly different proliferation activity.

Under normal physiological conditions, once G1/S transcription is fully activated the cell cycle most often completes (Donjerkovic et Scott 2000). Consequently, the G1 phase represents a time window during which a cell can opt for other cell fate choices like quiescence (Sagot et Laporte 2019; Cheung et Rando 2013) or differentiation (Grinenko et al. 2018), and extending the duration of G1 phase increases the odds that other developmental decisions are made at the molecular level (Burda et Roeder 2022; Calegari et Huttner 2003). How are cell division, quiescence and differentiation coordinated by the G1/S network? This is the central question motivating this work.

To study how cell fate decisions are coordinated, the hematopoietic system is an ideal model: each day, the bone marrow produces ∼284 billion new hematopoietic cells, accounting for ∼86% of total cellular turnover in the human body (Carlberg et Velleuer 2022). This output originates from a small population of hematopoietic stem cells (HSCs). To sustain the lifelong hematopoiesis, most HSCs are dormant in cell-cycle quiescence (G0 phase), i.e., have a low proliferation rate that protects them from functional exhaustion and cellular insults (Nakamura-Ishizu, Takizawa, et Suda 2014). A subset of HSCs, however, differentiates into multipotent progenitors (MPPs), which further give rise to lineage-committed progenitors (X. Zhang et al. 2024). Progenitor populations are characterized by a fast proliferation (Matsumoto et Nakayama 2013), which supports self-renewing of progenitors population and the maintenance of stem cell identity (Ruiz et al. 2011; Galloway 2024). Upon response to differentiation cues, lineage-committed progenitors evolve towards terminally differentiated cells that display a low intrinsic proliferation rate, although some cell types such as naïve and memory B an T cells, can re-enter the cell cycle upon contact with an antigen (Muroyama et wherry 2021). Hence, cell cycle activity evolves along the hematopoietic differentiation path, raising the question of the functional role of cell cycle dynamics in the acquisition of differentiation biases towards lineage commitment, and vice-versa.

Before committing to further differentiation, cells sense differentiation cues and make cell fate choices during G1 (Dalton 2015), the duration of which varies depending on the cell type and environmental context (Mende et al. 2015). Increased G1 duration delays cell cycle progression increases cells’ exposure to differentiation cues. In contrast, a short G1 drives the rapid proliferation of progenitors and uncouples cells from extrinsic signals. Thus, the G1/S checkpoint not only serves as a gate to proliferation but also constrains alternative fate choices. Interestingly, changes in cell cycle duration alone can shift developmental trajectories (Dalton 2015). For instance, increased cell cycle speed in megakaryocytic-erythroid progenitors (MEPs) is sufficient to promote their differentiation towards erythroid progenitors (ERPs) rather than towards megakaryocytic progenitors (MkP) (Lu et al. 2018a), Similarly, a slower cell cycle in fetal liver progenitors promotes myelomonocytic over B cell fate (Kueh et al. 2013). Thus, cell cycle duration, and G1 duration in particular, regulates lineage specification.

Across kingdoms of life, passage through G1/S is orchestrated at the transcriptional level by the E2F family of transcription factors: in plants, E2Fa-c, reviewed in (Gombos et al. 2023); in yeast, SBF/MBF, reviewed in (Hendler et al. 2018) across evolution; and in mammals, E2F1-3, reviewed in (Attwooll, Lazzerini Denchi, et Helin 2004). In G1, E2Fs are kept inactive through high affinity binding of Retinoblastoma (RB) family proteins (Rbr in plants, whi5 in yeast, and RB1/ RBL1-2 in mammals, (weinberg 1995; Henley et Dick 2012; Chinnam et Goodrich 2011; Gombos et al. 2023)). In late G1, RB-like proteins are heavily phosphorylated by cyclin dependent kinases (CDKs, CDK2-4-6 in mammals). The intrinsic kinase activity of endogenous CDKs is very low but is boosted upon binding to G1 cyclins (Cln1-3 in yeast, cyclins D1-3 and E1-2 in mammals (Malumbres 2014)). RB phosphorylation disrupt RB-E2F complexes, releasing active E2Fs that transcribe genes required for S-phase progression. Among E2F targets are the *CCNE1* and *CCNE2* (cyclin E1 and E2), *CDK2* (cyclin-dependent kinase 2) and *CDC25A* (cell division cycle 25A) genes, that further activate CDKs and complete RB phosphorylation in a positive feedback loop (Skotheim et al. 2008; weinberg 1995; Ohtani, DeGregori, et Nevins 1995)). In human, the activity of Cyclin-CDK complexes is balanced by cyclin-dependent kinase inhibitors (encoded by *CDKN* genes). p16 (*CDKN2A*) and p15 (*CDKN2B*) specifically inhibit CDK4/6, while p21 (*CDKN1A*), p27 (*CDKN1B*) and in certain tissue contexts p57 (*CDKN1C*) also inhibit CDK2 and may also play an activatory role by assisting in the assembly of CycD-CDK4/6 complexes. Dysfunctional *CDKN2A/B* is the 2^nd^ most common genetic alteration found in cancer, and is also very frequent in hematological malignancies (Teierle et al. 2023).

The consensus chain of events that lead to full G1/S commitment is the following: first mitogenic signals and/or growth factors induce the transcription of cyclin D1-3 genes (Sewing et al. 1993; Roussel et al. 1995; Surmacz et al. 1992; Matsushime et al. 1991), promoting assembly of CycD-CDK4/6 complexes that initiate RB phosphorylation, partially reducing E2F inhibition which induces low-level transcription of cyclin E and other activators, activating the positive feedback loop that leads to full RB phosphorylation, E2F release and G1/S activation via transcription of E2F targets. In this paradigm, absence of either CycD-CDK4/6 or CycE-CDK2 would prevent full RB phosphorylation and E2F activation (see (Hume, Dianov, et Ramadan 2020) and discussion therein), keeping E2F in an intermediate state of moderate activity (Konagaya et al. 2024), insufficient to pass G1/S. However, some cell type proliferates despite minimal expression of CycD-CDK4/6 related genes, possibly because RB hyperphosphorylation is already achieved at cell birth in some contexts (Spencer et al. 2013). Likewise, the existence of cancer cells resistant to CDK4/6 inhibition challenge this canonical model, and demonstrates the existence of alternative routes through G1/S (Knudsen, witkiewicz, et Rubin 2024; Kim et al. 2023).

we hypothesize that distinct cell types make a differential use of the same G1/S molecular network to pass the transition. This idea is supported by the fact that while certain cell types degrade RB during G1 phase to elicit E2F activation (e.g. in hTERT-immortalized human mammary epithelium cells, (Zatulovskiy et al. 2020)), and depend on CDK activity, others pass G1/S without RB degradation (e.g. hTERT-immortalized human retina epithelial cells (Sanidas et al. 2019; Sun et al. 2017), indicating a plethora of strategies cells can employ to activate E2F and commit to division. we hypothesize that, due to the large variability in gene expression patterns and proliferation rates across hematopoietic cell types, the hematopoietic system encloses the different transition dynamics encoded within the G1/S network.

To validate this hypothesis, we leveraged the best-resolved transcriptomics atlas of healthy human hematopoiesis published to date (X. Zhang et al. 2024), which provides a detailed snapshot of the relative expression of core G1/S regulators across different cell types. we integrated this data into a dynamical mathematical model of the G1/S transition, that enabled to predict the proliferative potential of each cell type and provided a mechanistic basis into how different cell types navigate the G1/S transition. Our analysis identified 3 major routes for the G1/S transition, and the hematopoietic cell types that elicit each route. Since the different routes through G1/S are based on distinct G1/S core regulatory proteins, our model explains the differential sensitivity of hematopoietic cell types to specific drugs (e.g. CDK4/6 inhibitors). we envision our model could serve as a basis for personalized therapeutic strategies and foster the clinical investigation of novel targeted therapies and rational drug combinations (Bride et al. 2021).

## Material and methods

### G1/S genes

Throughout the manuscript, we consider a core subset of 24 genes/proteins that are the most direct regulators of the G1/S transition. The list of G1/S genes was assembled from various literature sources and include: the activating E2F transcription factors (*E2F1-3*); E2F repressors *RB1*, *RBL1* and *RBL2* genes; CDKs relevant to E2F de-repression, i.e. *CDK2/4/6,* and their activators cyclins D1-3 (*CCND1-3* genes) and cyclins E1-2 (*CCNE1-2*); 8 *CDKN* genes coding from CDK inhibitory proteins (*CDKN1A-C, CDN2A-D*, and *CDKN3);* and finally, the CDC25A phosphatase that activates CDK2-CycE complexes, and the c-MYC transcription factor that partially controls CDC25A and cyclin E expression, independent of E2F. The set of 24 genes listed above (3 activating *E2F*, 3 repressing *RB*-like, 3 *CDK*, 5 cyclins, 8 *CDKN*, *CDC25A* and *MYC*) are the basis of our mathematical model and are referred to as the “G1/S genes” in the transcriptomics data analysis detailed below and throughout the manuscript (Table 1).

**Table 1.** G1/S genes. List of G1/S regulatory genes accounted for in this study (core G1/S network, see Fig. 1).

### Single cell transcriptomics data curation

Healthy adult bone marrow single cell RNA sequencing (scRNA-seq) data (Hay et al. 2018) was downloaded from synapse (syn18659336) as a h5ad object, and updated with cell type annotation from (X. Zhang et al. 2024) downloaded from syn53237600. The data was processed using the toolkits Scanpy (v1.10.4) (wolf, Angerer, et Theis 2018) in Python and Seurat (v.4.4.0) (Butler et al. 2018), with <u>anndata</u> (v.0.7.5.6) in R (v.4.3.1). Gene annotation was updated based on ENSG IDs to Ensembl version 113 (release October 2024) (Dyer et al. 2025) via Biomart in R (v.2.58.0). In case of duplicated gene identifiers, the version with the higher gene count and lower Entrez gene ID were kept.

### Data processing

we used the finest cell type annotation provided by Zhang and co-authors (“Level 3”, (X. Zhang et al. 2024)) to process the single cell data and create pseudobulk samples. To focus on high-quality hematopoietic cells, we filtered out stromal cells and excluded cells with a mitochondrial read fraction >10% (1117 cells) as recommended in (Osorio et Cai 2021). Moreover, we removed outlier cells with a log-library size or log-transformed number of expressed genes exceeding >5 median absolute deviations from the median of the given cell type. Then we aggregated read counts from cells from the same donor-cell type combination with Decoupler (version 1.8.0) to form pseudobulk samples. Samples formed from < 10 cells or < 50,000 read were removed as recommended (Robinson, McCarthy, et Smyth 2010). Furthermore, cell types present in <5 donors were excluded from the dataset. This procedure yielded 68 cell types, totalizing 95,685 single cells and 501 pseudobulk samples (Supplementary Table T1). For the coarse level pseudobulk analysis, we utilized “Level 1” annotation of the same dataset provided by Zhang et al. to form pseudobulk samples. For Principal Component Analysis (PCA) analysis, we further reduced the number of cell types to 24, i.e., the number of core G1/S genes, by excluding the cell types with the lowest number of donors (Ba/Ma/ Eo).

### Data normalization and RNA concentration estimate

To normalize pseudobulk samples for both the coarse (Level 1) and fine (Level 3) annotation levels, library sizes (i.e., total RNA count per pseudobulk sample) were adjusted by the TMM method, and counts per million (CPM) and log2-CPM counts were calculated for each gene using the cpm (log= FALSE or log=TRUE) function in EdgeR (v.4.0.16) (Yunshun Chen et al. 2025).

To estimate average RNA concentration per gene and cell type, for each gene G and donor-cell type sample S, we calculated the average of single cell-level normalized expression values (i.e., gene UMI counts divided by total UMI per cell, scaled to 10,000):

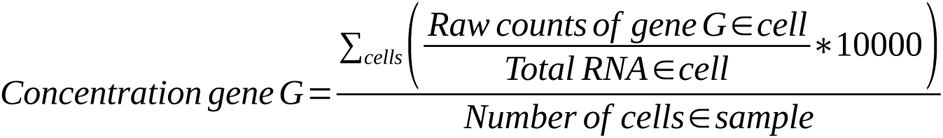

To calculate average concentration of E2F (E2F1+E2F2+E2F3), CDK (CDK2+CDK4+CDK6), RB (RB1+RBL1+RBL2), and CCNE (CCNE1+CCNE2) (Figs. 6 and 7), raw UMI counts of individual genes (e.g. E2F1+E2F2+E2F3) were summed per cell prior to normalization. For model input, concentrations of the 24 G1/S genes were averaged across donors to obtain one value per cell type (Supplementary table T8).

Average cellular transcript abundance of E2F per cell type was calculated by summing raw UMI counts of E2F1+E2F2+E2F3, divided by the number of cells per individual donor-cell type combination (i.e., “absolute scaling” in (Kim et al. 2023)).

Conclusions presented in the manuscript were all based on the pseudobulk samples, or fractions of those pseudobulk samples (Fig. 8). However, for visualization purpose, we also introduced graphs where gene expression at the single cell level are presented.

**Figure 1.**
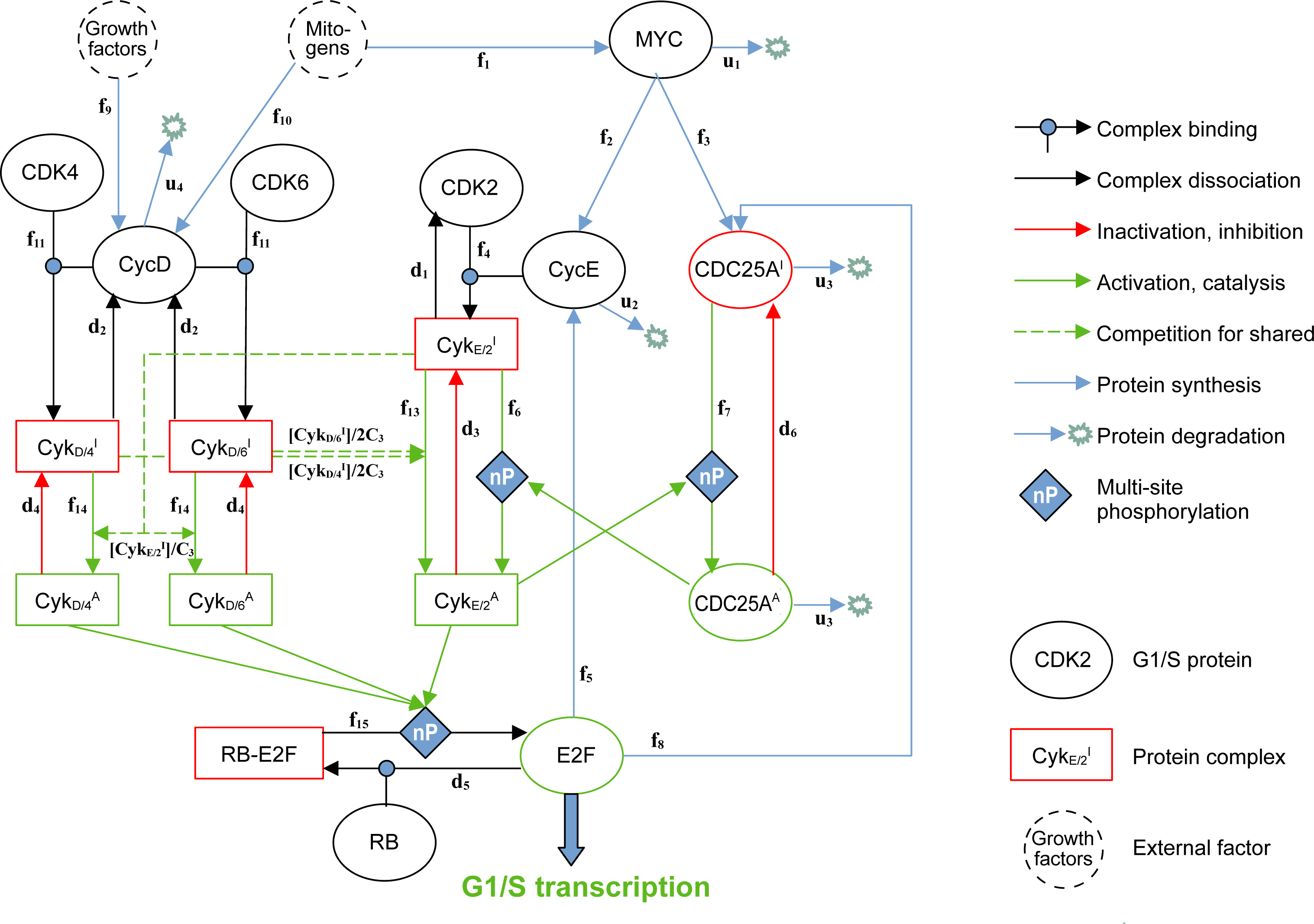
A mathematical model for the G1/S transition. Schematics of the G1/S network model. Molecular species and complexes an are shown as ellipses and rectangles respectively, framed in green and red depending on whether they are in active or inactive form. Dashed circles represent upstream mitogenic inputs that influence commitment to cell-cycle entry. Black ellipses represent proteins whose activity towards the G1/S transition is not accounted for in the model, when they are not part of complexes. Arrows indicate all biochemical reactions included in the model, with the corre sponding rate/coefficient shown on the side of the arrow. Blue arrows indicate synthesis/degradation reactions, black arrows complex binding/dissociation, red arrows complex/protein inactivation and green arrows complex/protein activation. Dashed green arrows represent indirect activation that stem from the competition for shared CDK inhibitor proteins, and modulate the rates f13 and f14. “nP” diamonds represent activation functions that are governed by multi-site phosphorylation, multiplying the corresponding rate by a Hill function of the activatory factor (see Methods, Supplementary Methods).

**Figure 2.**
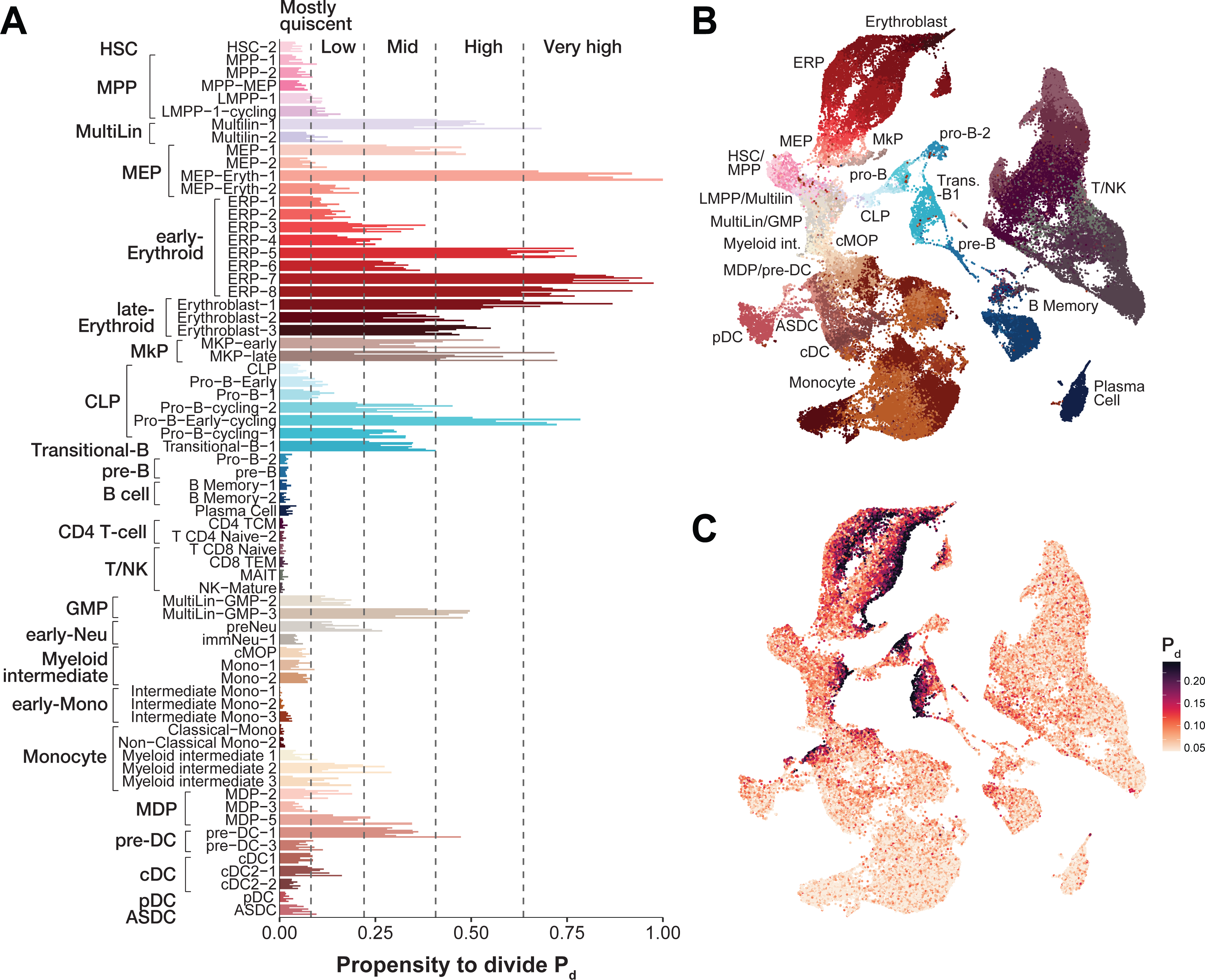
The propensity to divide evolves non-monotonically along branches of the hematopoietic tree, consistently across donors. A) Bar chart shows the propensity to divide score (P_d_) in developmentally ordered hematopoietic cell types from the adult bone marrow ((Hay et al. 2018), scRNA-seq annotated by (X. Zhang et al. 2024)). For each cell type, individual bars represent the donors, highlighting low inter-donor variability in P_d_. Coarse cell type annotations (Level 1, (X. Zhang et al. 2024)) are indicated in bold. Dashed lines indicate P_d_ categories (Jenks breaks). B) UMAP representation of the dataset at single cell resolution (1 dot = 1 cell, see Methods) captures hematopoietic lineage relationships and highlights differentiation trajectories. Cell types are colorcoded as in A). C) UMAP (panel B) colored by P_d_ scores illustrates how the proliferative potential evolves during hematopoiesis.

**Figure 3.**
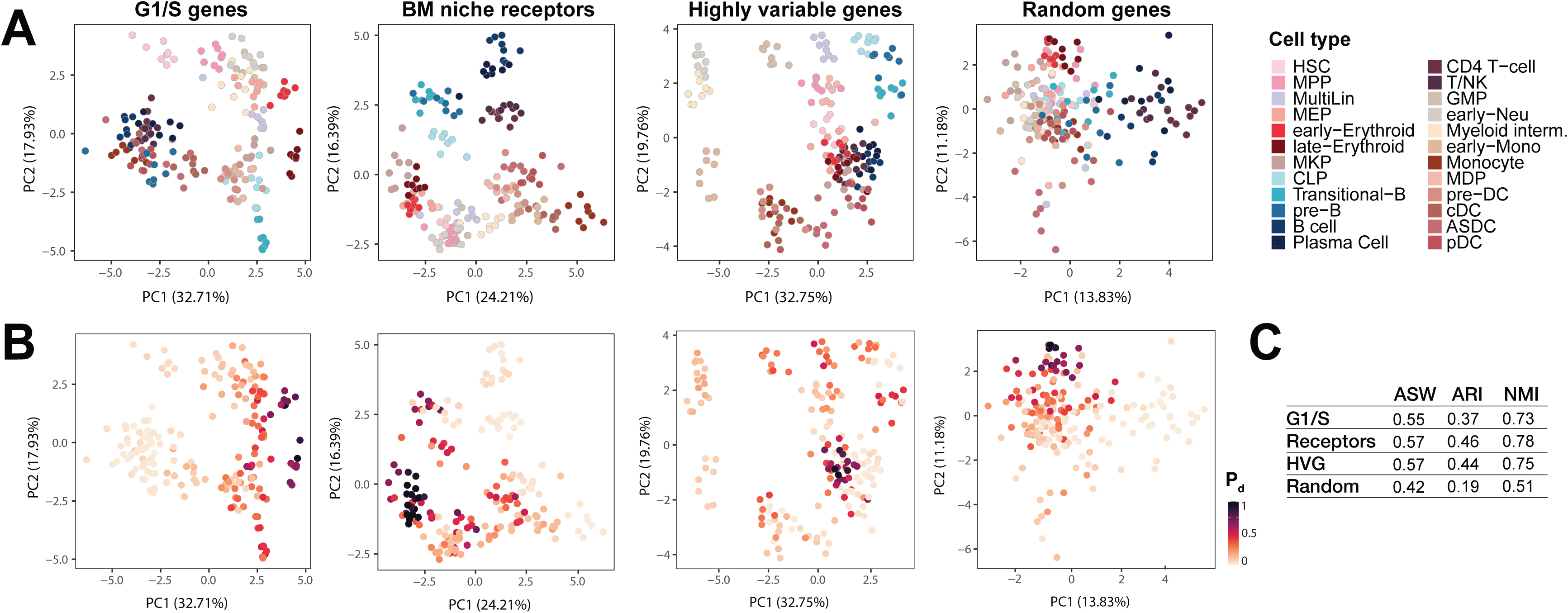
The expression of 24 core G1/S genes accurately segregates hematopoietic cell types and captures their propensity to divide. A) PCA of pseudobulk samples (*i.e*., aggregated cells from each cell type and donor) based on the expression of 24 G1/S (Table 1), receptors for bone marrow (BM) niche signals (Supplementary Table T3), highly variable genes (HVG) and random genes. Each dot represents one donor (n=8). Clustering performance for each 24-gene set is shown. Same PCA plots as in A) recolored according to the propensity to divide score P_d_ of each pseudobulk sample.

**Figure 4.**
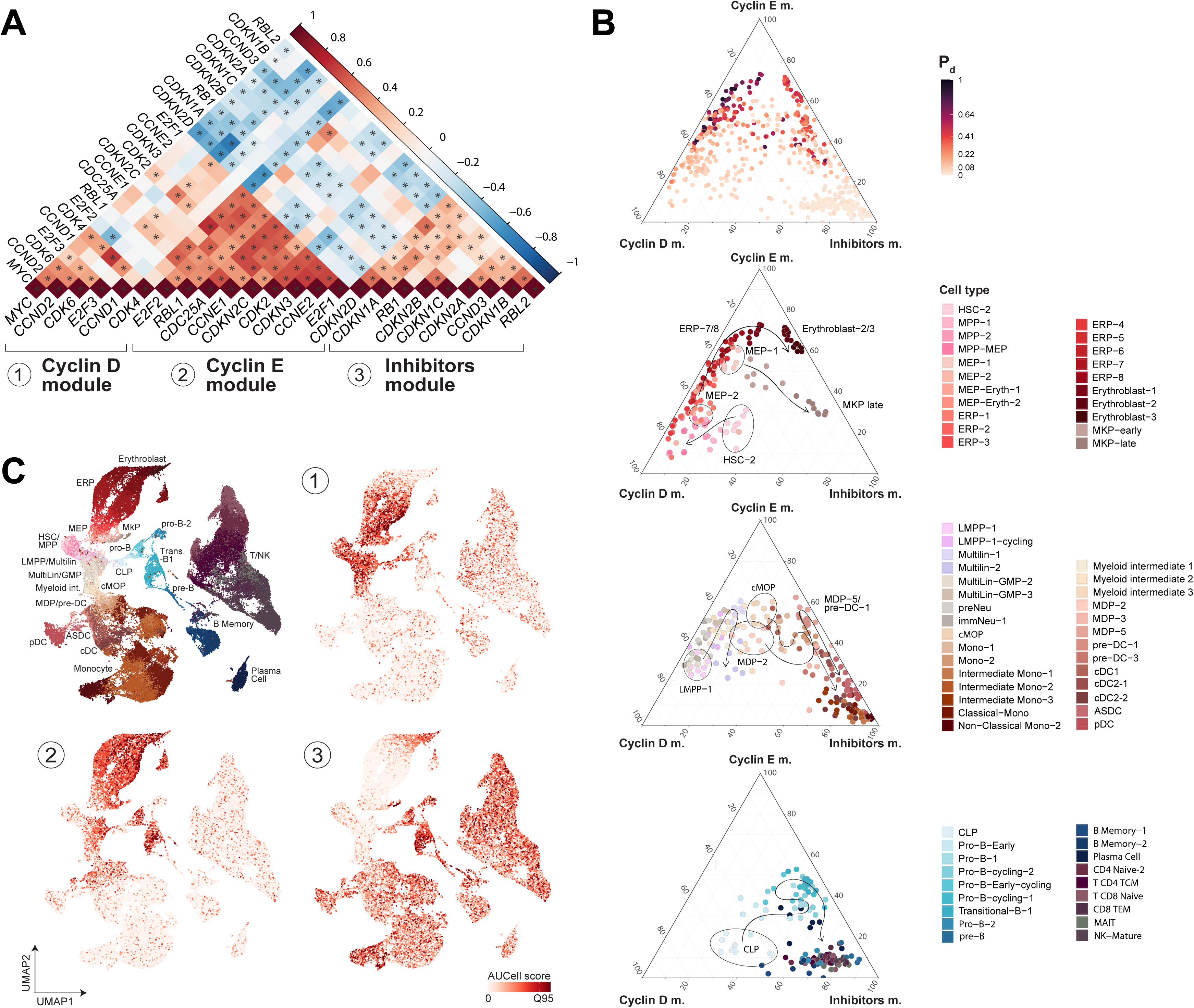
Three distinct G1/S network sub-modules are sequentially activated during hematopoietic differentiation. A) Hierarchical clustering of G1/S gene-gene Pearson correlations across all cell types (n= 8 donors, pseudobulk) identifies three distinct co-regulated gene groups: Cyclin D, Cyclin E and Inhibitors module. B) Ternary plot shows pseudobulk samples positioned according to their relative expression of Cyclin D, Cyclin E and Inhibitor module (one dot = aggregated cells from one cell type and donor). Top: cell types colored by their propensity to divide P_d_; middle top, middle bottom and bottom: lineage-specific G1/S module activity carves out a landscape through which hematopoietic cells traverse as they differentiate. Arrows illustrate differentiation trajectories. C) Single-cell UMAP representation of the dataset (from Fig. 2) highlights the sequential activation of Cyclin D, Cyclin E and Inhibitors module along the hematopoietic tree.

**Figure 5.**
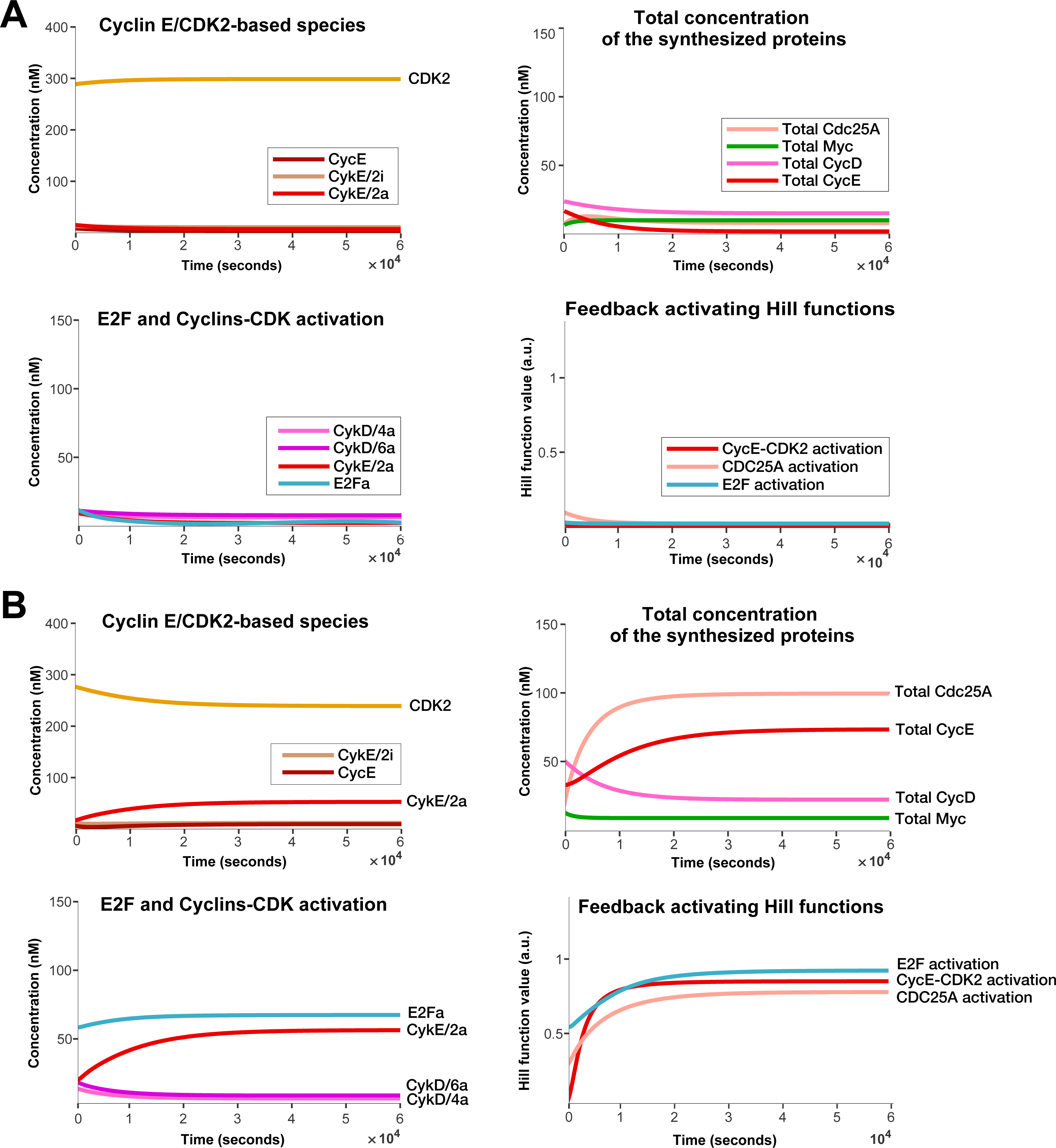
Our mathematical model predicts a bistable G1/S transition. Each panel shows the simulated time course (x axis, in seconds) in protein concentration changes of the core G1/S protein/protein complexes (y axis, concentration in nM) or Hill function activation (in arbitrary units). Time courses were simulated under different initial conditions, to produce convergence towards the low activity steady state (A) or the high activity steady state (B). All timecourses were simulated under the default parametrization (Table 2).

**Figure 6.**
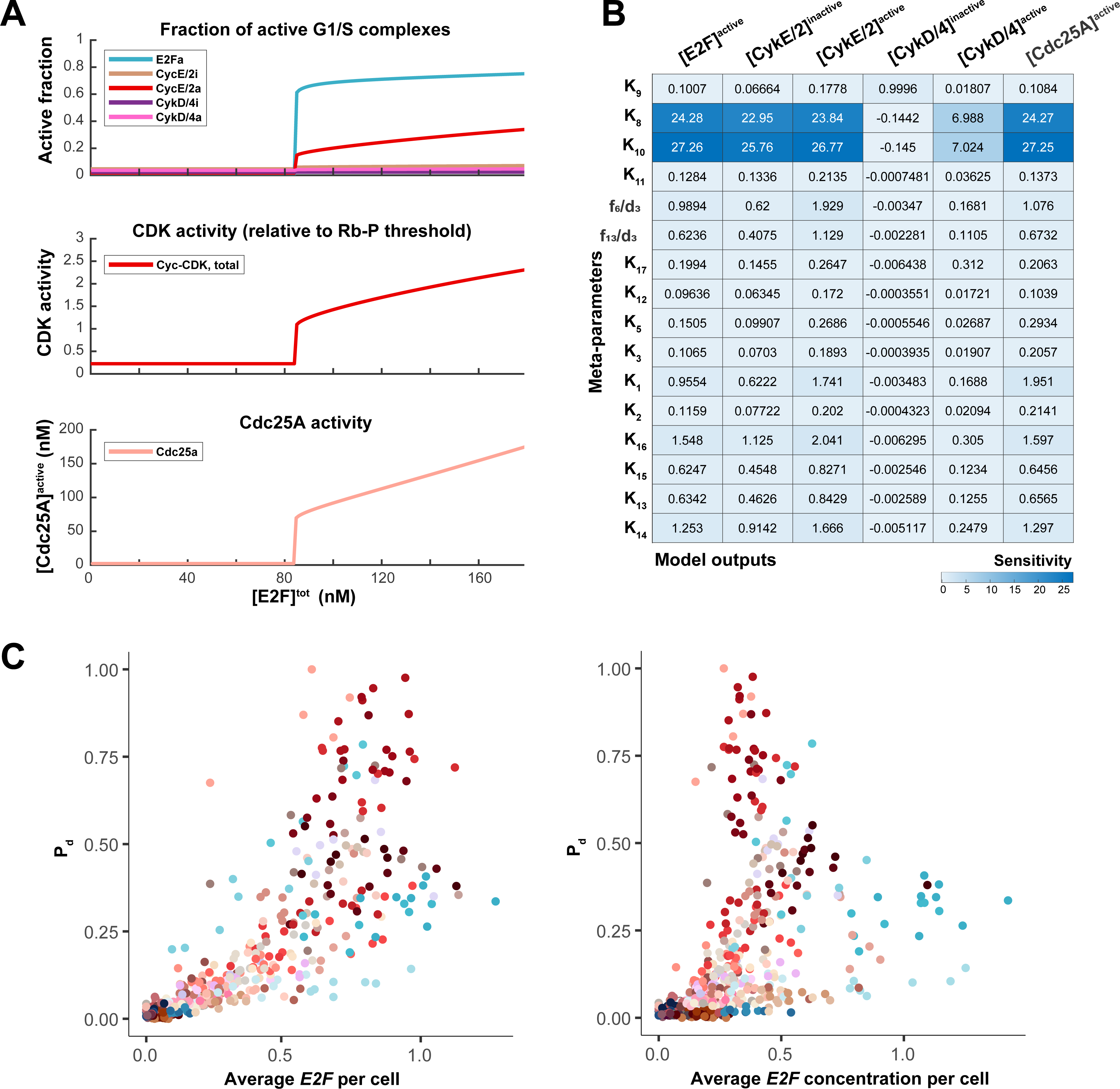
E2F accumulation leads to a sharp G1/S transition, triggered by a dynamical instability. **A)** Steady state concentrations (y-axis, in nM) or fraction of activated complexes (as indicated) as a function of total E2F concentration (x-axis, in nM). Default total RB concentration is 100 nM. B) Sensitivity analysis of the steady-state concentrations (model outputs, in columns) in response to a 1 % change of the meta-parameters (rows), normalized to this change (see Methods), at the tipping point (in A) where [E2F]tot = 85 nM. C) Propensity to divide P_d_as a function of the average E2F mRNA abundance per cell (left) or of the average concentration of E2F mRNAs per cell, computed as the average single-cell concentration per cell type for each donor (Methods). Color-coded according to cell types.

**Figure 7.**
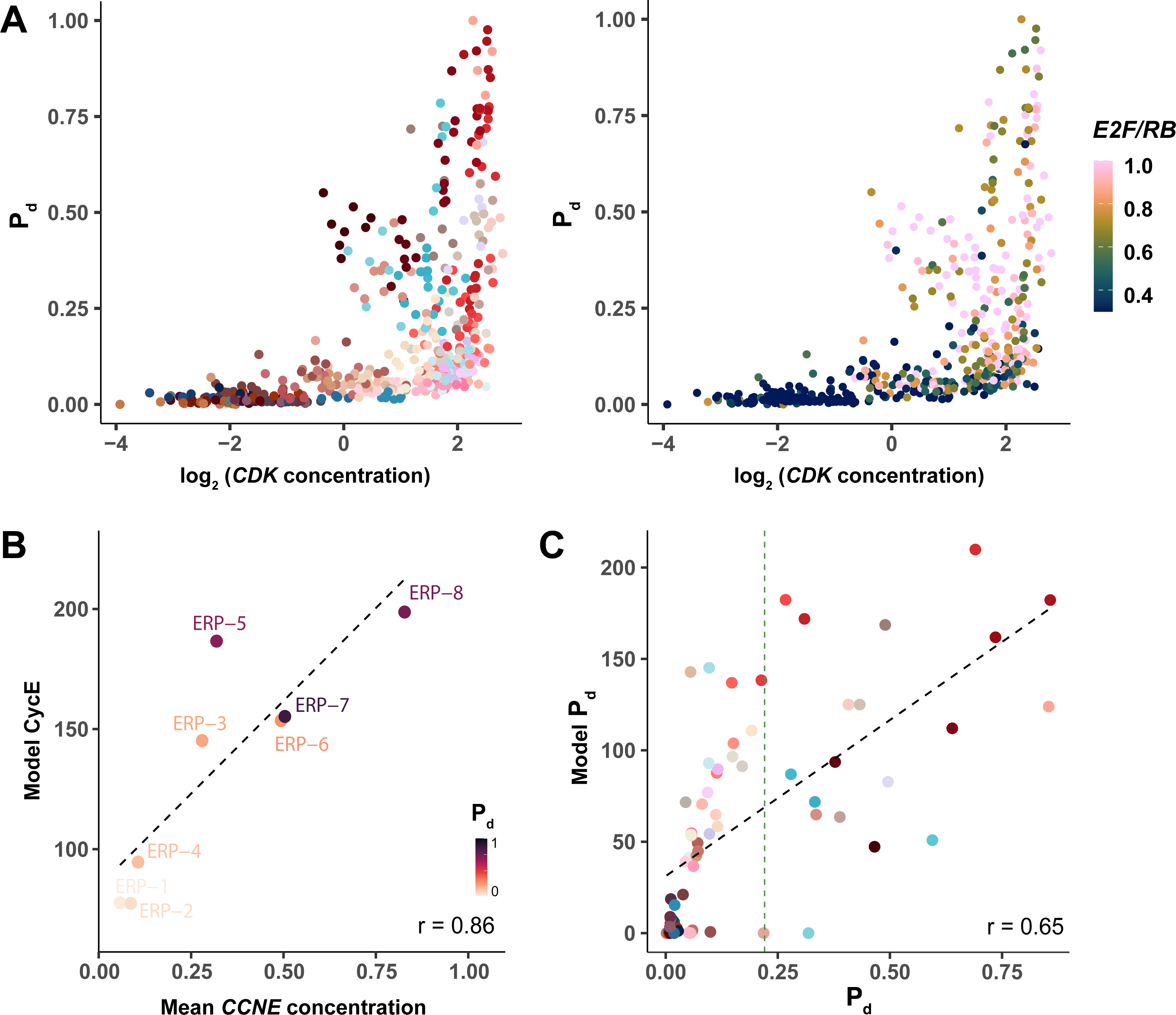
Pro-B cells, late MDP/pre-DC cells, and Erythroblasts opt for E2F-driven proliferation whereas MEP/ERP cells rely on CDK-driven proliferation. A) Propensity to divide P_d_ (yaxis) as a function of CDK mRNA concentration (x-axis), color-coded by cell type (left) or by the total E2F/RB ratio (right). B) Total CycE protein concentration predicted by the model (y-axis, nM) as a function of the mean *CCNE1/2* transcript concentration (normalized UMI counts) in erythroid progenitor developmental stages (y-axis), demonstrating a strong correlation. Color-coded by average P_d_. C) Average propensity to divide P_d,model_ predicted by the model (y-axis) versus P_d_ estimated from scRNA-seq profiles (x-axis), across 68 hematopoietic cell types (color coded). Vertical green line indicates the limit P_d_ for mostly quiescent and low proliferation categories (Fig. 2, Jenks breaks).

**Figure 8.**
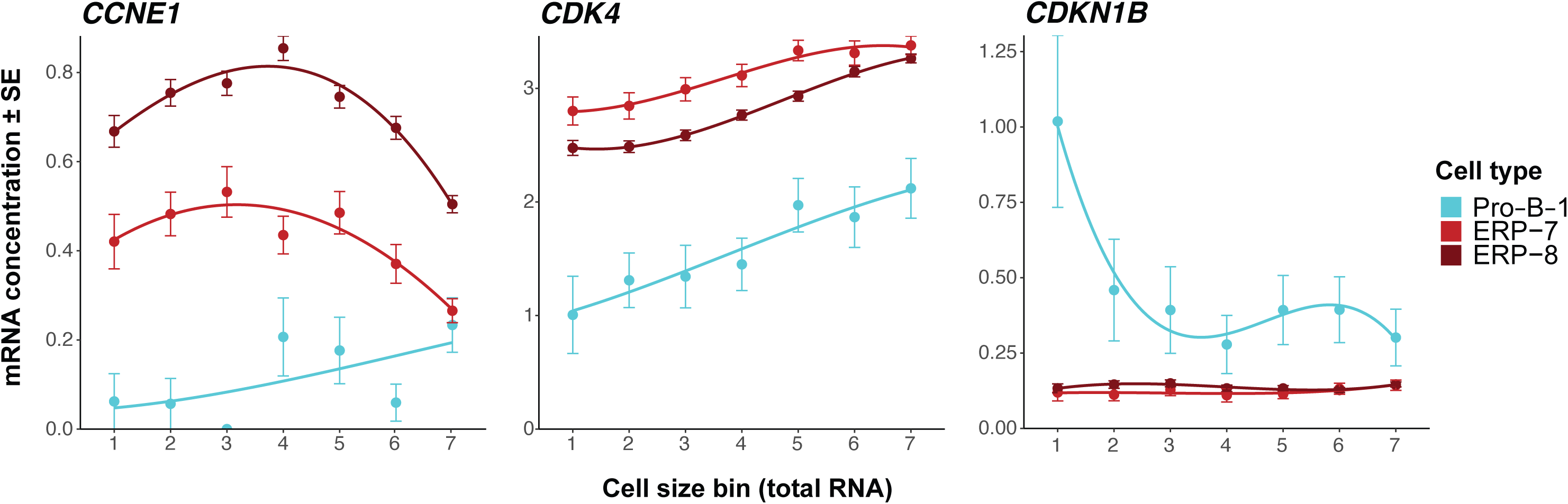
Cell type-specific differential scaling of G1/S regulators during G1 growth elicit distinct transition mechanisms in B lymphoid progenitors and late ERP. Single cell gene concentration averaged over all cells of a given type across donors (y axis) as a function of their total RNA content (proxy for cell size, x axis), color-coded according to cell types. Curves represent 3rd- order polynomial fits.

### Single cell level visualization tools

To visualize the structure of the hematopoietic transcriptome dataset, we generated a UMAP (Uniform Manifold Approximation and Projection) representation of the single cell data following a standard processing pipeline for dimensionality reduction: first, gene expression was normalized to the cell’s total RNA counts (i.e., library size) and multiplied by 10,000, followed by log1p transformation via the function NormalizeData, and finally z-scored with ScaleData in Seurat. Then, the top 4000 genes with highest feature importance were detected by a Random Forest classifier (implemented in Delve benchmark (Ranek et al. 2024), and used to compute principal components (PCs) via the RunPCA function. Finally, the top 30 PCs were used to produce the UMAP plot using RunUMAP (method = uwot) in Seurat. For a possibly faithful representation of the data, the hyperparameters of the UMAP were optimized with scDEED package (v.0.1.0)(Xia, Lee, et Li 2024). The resulting UMAP (Fig. 2B) where our 68 hematopoietic cell types align along developmental trajectories, was used as a backbone for overlaying various scores (see below), providing an overview of how the different scores vary along developmental trajectories across the hematopoietic tree.

### Propensity to divide from transcriptomics data

The propensity to divide P_d_ was defined based on the mRNA counts of genes that are expressed specifically after the G1/S transition, *i.e*., during the S, G2, and M phases of the cell cycle. S/G2/M genes were obtained from Macosko et al. (clusters 6-7-8, 277 genes in total, (Macosko et al. 2015)). This list was intersected with the list of universally essential genes, i.e. genes whose knock-out suppresses viability across 10 different cell lines (Bertomeu et al. 2018), to derive to a list of 29 core S/G/2M genes whose expression specifically after the G1/S transition is required across all viable cell types (Supplementary Table T2).

At the pseudobulk (cell type-donor combination) level, P_d_ was calculated as follows:

1. For each S/G2/M-specific gene *G*, and each pseudobulk sample S, we calculated the mean *G* expression per cell:

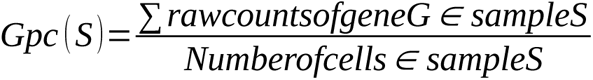

2. To balance each gene’s *G* contribution to the score, we z-scored each *G* mean expression across samples:

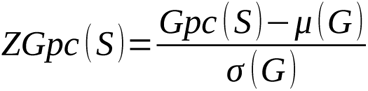

where μ(G) and σ(G) are the mean and standard deviation of gene *G*’s expression across all samples.

3. we then calculated the average z-scores across all the genes *G* of the S/G2/M gene list, for each pseudobulk sample *S*:

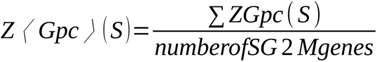

4. Finally, the average Z-scores of all samples were rescaled to the [0-1] interval via:

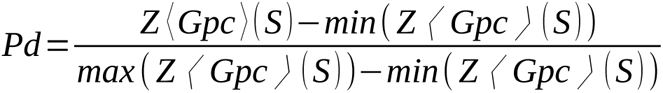

where *min*( *Z* ⟨ *Gpc* ⟩ ( *S* ) ) and *max* ( *Z* ⟨ *Gpc* ⟩ ( *S* ) ) are respectively the smallest and largest average Z-score across all pseudobulk samples. P_d_ values were segregated into 5 categories using Jenks natural breaks implemented in classInt (v.0.4.11, see Fig. 2A).

At the single cell level, for visualization purposes, P_d_ was estimated using the AUCell algorithm (v.1.22.0), which uses a ranking-based approach to calculate the enrichment of a gene list relative to other genes expressed in the same cell (Aibar et al. 2017). The list of 29 S/G2/M specific genes was used as an input to AUCell.

### Principal Component Analysis, correlation analysis and definition of Cyclin D, Cyclin E and Inhibitors modules

Principal Component Analysis (PCA) was performed using z-scored log2-CPM counts obtained from coarse-level (level 1) pseudobulk samples. P_d_ was calculated for those samples as described above for the fine-level analysis, and overlaid on the PCA plots. To evaluate how well different cell types separate in the PCA space, we utilized clustering performance metrics Adjusted Silhouette width (ASw), Adjusted Rand Index (ARI) and (Normalized Mutual Information) NMI (Luecken et al. 2022) in scib (v.1.1.7).

Gene-gene correlograms were plotted from the Pearson correlation coefficients of z-scored log2-CPM counts between any pair of genes, across all the 68 level 3 pseudobulk samples, using the corrplot package (v.0.95, (wei et Simko 2024)). Genes were ordered using hierarchical clustering. The statistical significance of gene-gene correlations across the pseudobulk samples was assessed with the cor.mtest function, and a confidence level of 0.99. we adjusted the p-values for multiple hypothesis testing using the Benjamini-Hochberg correction method, and considered correlations with FDR<0.01 as significant. Correlogram highlighted different patterns of correlation for cyclin D related genes, cyclin E related genes and G1/S inhibitor genes, from which we derived 3 modules (see Results). For a better visualization of the shared correlations, *CDK4* has been swapped with *CDKN2C* to highlight its unique position, as *CDK4* correlated strongly both with Cyclin D and Cyclin E modules (Supplementary Table T4).

To position pseudobullk samples according to how they express the three modules, we used barycentric coordinates. Specifically, for each sample (donor-cell type), TMM-adjusted CPM counts per module were z-scored, averaged, and min-max scaled as done in steps 2-4 for P_d_ calculation. Finally, scaled module scores were divided by the sum of the 3 scores for the same sample, ensuring that once re-normalized, Cyclin D module score + Cyclin E module score + Inhibitors module = 1, prior to conversion to barycentric coordinates and ternary plotting using ggtern (v.3.5.0) package. To visualize the overall expression of cyclin D, cyclin E and Inhibitors modules at the single cell level, we scored the enrichment of each module using the AUCell tool, as described above for the propensity to divide. Heatmaps were generated with ComplexHeatmap (v.2.18.0)(Gu 2022) with z-scored log2CPM expression values clipped to ∣z∣ ≤ 2 to prevent extreme values skewing the visualization. Cus tom code is available at: https://github.com/andreahanel/2025

### Interactions amongst the G1/S regulatory proteins: construction of the G1/S molecular network

Interactions between the 24 core G1/S genes defined above were pulled from the literature. The mechanisms of action of *CDKN* gene products differ in that CDK4/6 specific inhibitors (INK4 family, *CDKN2* genes) alter the structure of the Cyclin-binding pocket on CDK4/6, preventing the formation of the CycD-CDK4/6 complexes specifically, while *CDKNs* of the Cip/Kip family (*CDKN1* gene products) can bind to an already formed Cyc-CDK complex, rather inhibiting its catalytic activity (Jeffrey, Tong, et Pavletich 2000). The mechanism of CycE-CDK2 inhibition by *CDKN3* is less clear. The expression of E-type cyclins is under control of E2F, producing a positive feedback loop where a small pool of active E2F transcribed *CCNE1-2* genes, increasing CycE-CDK2 activity which phosphorylates more RB-like proteins, increasing the overall level of E2F activity. In addition to cyclin E, E2F also induces expression of CDC25A (Vigo et al. 1999), a phosphatase with documented activating effect on CDK2 (Donjerkovic et Scott 2000). The phosphatase function of CDC25A is activated by CycE-CDK2 (Hoffmann, Draetta, et Karsenti 1994), establishing another positive feedback loop within the G1/S network. we note that the *CDC25A* gene is a transcriptional target of c-MYC independent of E2F (Galaktionov, Chen, et Beach 1996), making c-MYC a core element of the G1/S network. c-MYC expression is, itself, responsive to mitogenic signals (Miller et al. 2012), while D-type cyclins expression also responds to a range of growth factors (Sewing et al. 1993; Roussel et al. 1995; Surmacz et al. 1992; Sherr 1996; Matsushime et al. 1991), coupling the cell cycle trigger to the environment.

### Simplification of the G1/S network and mathematical description

To simplify the network and its mathematical description, we chose to represent explicitly E2F1-3 as a single E2F factor, RB-like proteins as a single RB transcriptional repressor, cyclin E1-2 as a single CycE, cyclin D1-3 as a single CycD, and to include the effect of CDK inhibitor proteins in the model parameters describing the binding of CycD to CDK4/6 (for *CDKN2* gene products) and describing the activation of CycD-CDK4/6 and CycE-CDK2 complexes (for *CDKN1* gene products). while *CDKN1A,C* inhibits only CycE-CDK2, *CDKN1B* can also impact CycD-CDK4/6, prompting us to include potential differential inhibition of CycE-CDK2 and CycD-CDK4/6 complexes by *CDKN.* Therefore, the corresponding parameters in the model can vary independent of each other. In summary, our model includes explicitly 9 protein factors: E2F, RB, CycD, CycE, CDK2, CDK4, CDK6, CDC25A and (c-)MYC, that represent 16 proteins of the core G1/S network, and includes implicitly 8 CDK inhibitor proteins whose effects are captured by changes in model kinetic parameters d_3_ and d_4_ (see below and Supplementary Methods).

Total concentrations of proteins with long half-lives (CDK2, CDK4, CDK6, E2F, RB, several hours, (Mathieson et al. 2018)) were assumed constant and considered as model parameters, i.e. external inputs to the G1/S network. Binding/unbinding reactions were assumed to satisfy mass-action kinetics. Moreover, we chose to describe explicitly active and inactive forms of the complexes. Under those constraints, the full dynamics of the CDK2, CDK4, CDK6, E2F and RB sub-network is captured by 10 ordinary differential equations (ODEs) for the concentrations of 10 molecular species: unbound (inactive) CDK2-4-6 ([CDK2-4-6]^I^), cyclin-bound but inactive CycE-CDK2 (shortened as [Cyk_E/2_]^I^) and CycD-CDK4/6 (shortened as [Cyk_D/4-6_]^I^), cyclin-bound and catalytically active CycE-CDK2 (shortened as [Cyk_E/2_]^A^) and CycD-CDK4/6 (shortened as [Cyk_D/4-6_]^A^), and finally RB-free E2F ([E2F]^F^). Those 10 ODEs are provided as Supplementary Methods.

In contrast, total concentrations of CycD, CycE, CDC25A and (c-)MYC were assumed to be time-dependent variables, and their synthesis/degradation rates were also included in the model, in addition to their binding/dissociation to/from other factors, and their activation/inactivation. CycD synthesis was put under control of mitogens and growth factors; MYC synthesis was assumed to respond to mitogens; CycE and CDC25A synthesis was assumed to be stimulated by both active (RB-free) E2F and MYC (Zhou et al. 2020; Robson et al. 2011). Degradation rates were deduced from proteins half-lives as measured with SILAC proteomics (Mathieson et al. 2018) or biochemical experiments (CDC25A: (Shreeram, Hee, et Bulavin 2008), Myc: (Ahmadi et al. 2021; Thomas et Tansey 2011)). Cyclins were degraded only in the free state, but not while bound to CDKs, in line with the observation that subunits of protein complexes tend to be degraded at similar rates (the degradation rate of the Cyc-CDK complex (Mathieson et al. 2018; Masclef et al. 2019)). Hence, binding to CDKs stabilizes cyclins, as reflected by the very long CycE half-life in S-phase cells (Kossatz et al. 2010). Synthesis/degradation of CycD, CycE, CDC25A and (c-)MYC yielded 4 more ODEs, coupled to the binding/dissociation ODEs (Supplementary Methods).

Our model also describes explicitly the activation of CDC25A (via CycE-CDK2 phosphorylation (Hoffmann, Draetta, et Karsenti 1994)), activation of CDK2 (via CDC25A dephosphorylation at tyrosine 14 and threonine 15, (Hoffmann, Draetta, et Karsenti 1994)), and activation of E2F (via a Cyc-CDK multi-site phosphorylation-dependent dissociation rate of the RB-E2F complex), and the corresponding spontaneous deactivation reactions. The effect of multi-site phosphorylation on protein activity is complex, due to non-trivial conformational changes that follow each phosphorylation event and that may affect other putative phosphorylation sites. In this work, we consider the general framework of the Hill functions to model phosphorylation-dependent activation functions: 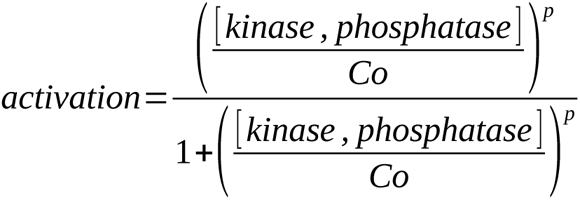, where the Hill threshold *Co* parametrizes the typical concentration range of the kinase/phosphatase that is required to elicit the activation, and the Hill exponent *p* parametrizes the sensitivity of the full activation to kinase/phosphatase concentration. *p* scales as the number of phosphorylation sites that need to be independently processed to elicit full activation (number of critical phosphosites (Dorsey et al. 2018)).

Finally, p27 (*CDKN1B* gene product) binds to and inhibits both CycE-CDK2 and CycD-CDK4/6 complexes (Sherr et Roberts 1999). Hence, Cyc-CDK complexes may compete for p27 and an increase in the concentration of inactive CycD-CDK4/6 (a fraction of which is p27-bound) leads to a decrease in the pool of available p27 and therefore an increase in CycE-CDK2 activation (Sherr et Roberts 1999). Our model accounts for this competition by using a [Cyk_D/4,6_]^I^ dependent activation rate of CycE-CDK2, and a [Cyk_E/2_]^I^ dependent activation rate of CycD-CDK4/6. All model equations are detailed in Supplementary Methods, and feature 15 variables and 38 parameters.

### Model parameters

Concentrations were expressed in nM. In the absence of proteome-wide measurements of absolute protein concentrations at the subcellular level in single human cells, we used different strategies to parametrize absolute total concentrations in our model. First, we used quantitative proteomics data (wiśniewski et al. 2014) to get an estimate of the order of magnitude of the G1/S proteins concentrations. In Ref. (wiśniewski et al. 2014), absolute protein counts are reported for various cell lines (A549, Hep-G2, PC-3 and U87MG) and range from about 3.000-5.000 copies of RBL1-2, 20.000-40.000 copies of RB1, 10.000-100.000 copies for CDK4 and CDK6, to 45.000-200.000 copies of CDK2 depending on cell types. Using an average volume of about 5 pL for those cell types we found the correspondence 30.000 copies ↔ 10 nM. Rieckmann and co-authors also measured RB1, RBL1-2, CDK2-4-6 copy numbers using spiked-in proteomics on primary human immune cells, in basal state and following immune activation (Rieckmann et al. 2017). They reported basal levels around 10.000-30.000 copies per cells depending on the proteins (with CDK2 and CDK6 more expressed than RBLs and CDK4) in most immune cells, and those number very multiplied several fold upon activation. Cyclin D2-3 and E2F3 expression was measured in just a few cell types to <10.000 copies per cell, indicating that total CycD levels at least 2-3 fold less than RB1. E2F1-2 were not reported. Of course, those values are averaged over a cell population and, importantly, over sub-cellular localizations, so that proteins with a much predominant nuclear localization (e.g. RB1) would show 5-10 fold larger nuclear concentration, since the nucleus is 5-10 times smaller than the cell. This rough analysis, though, informed that RB1 should be present at a typical concentration of 70-100 nM, 6-7 more than RBL1-2, and that CDKs should be presents at slightly larger concentrations (2-3 fold). Concentrations of MYC, Cyclins and CDC25A were not reported in quantitative proteomics datasets providing absolute values, possibly because of low expression levels. Those concentrations, while roughly estimated based on ad hoc conversion of quantitative proteomics data, were in reasonably good agreement with concentrations of homologs of these proteins in budding yeast, determined accurately using single live cell quantitative microscopy (Number and Brightness, N&B, (Dorsey et al. 2018)). In this work, the yeast RB1 (whi5) was quantified at 100 nM, and cyclins were expressed at 5 nM in early G1 and 5-15 nM in late G1 phase, at the transition. Our own N&B measurements in a NALM-6 pre-B leukemia cell model yielded a RB1 concentration of about 100nM and a very low p27 concentration (typically 10 nM, unpublished), indicating that p27 may be in stoichiometric default compared to Cyc-CDK complexes. Since no data was found for the absolute concentration of mammalian E2F1-3, we decided to keep the total concentration of E2F as a tunable parameter (see below, steady state analysis), and as a default value, to chose [E2F]^tot^=[RB1]^tot^=100 nM. Protein degradation rates were fixed to: CycD, 30 min, (Masclef et al. 2019); CDC25A: (Shreeram, Hee, et Bulavin 2008), Myc: (Ahmadi et al. 2021; Thomas et Tansey 2011); CycE: 30 min (Masclef et al. 2019). Synthesis rates were adjusted to provide steady state concentrations that fall in the range defined above.

For activation functions, Hill thresholds *Co*_1 ,2 ,3_ were chosen in the range of actual concentrations of the relevant proteins in the G1/S network (see steady state analysis), and the Hill exponents were chosen as the number of critical phosphorylation sites on the substrates RB1 ( *p*_3_=7), CDC25A ( *p*_2_=2) and CDK2 ( *p*_1_=1−5 as the number of critical CDK2 sites on CDC25A is unknown, default *p*_1_=3).

Cyc-CDK complex binding/unbinding dynamics were assumed to be in the millisecond (ms) range, to promote a fast activation of the targets when enough Cyc-CDK complexes are present. Binding rates were adjusted to yield ms dynamics even in the presence of low cyclin concentrations. The default parameters that were used unless otherwise specified are listed in Table 2.

**Table 2.** Model parameters and steady state meta-parameters. Default model parameter values (first tab), with rationales provided for the parameter choics. Meta-parameters that characterize the model’s steady state(s) are calculated in the second tab “meta-parameters” (rows 2-17) for 3 different values of the total E2F concentration (50 nM: lower than RB concentration; 100 nM: equal to RB concentration; 400 nM: 4 fold the total RB concentration). Rows 22-25: critical parameters for the bistability analysis (Supplementary Figs. S5-6), calculated for the default parameters (including a total E2F concentration of 100 nM) but for 3 different fractions of E2F activation (0%, 50%, 100%). The values shown for *K_6_, K_7_* and the two critical parameters 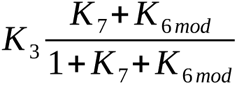 and 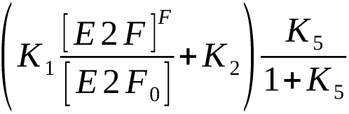 are to be used together with Supplementary Fig. S5 to visualize the model’s steady state(s).

### Model parameters for cell types other than default

The default model parameters above produced dynamics where the Cyclin E module dominates E2F activation. we matched those dynamics with that of pro-B cells, and grouped pro-B-early-cycling, pro-B-cycling-1, 2 and transitional pro-B1 to reduce inter cell type variability in the pro-B pool. For all the 24 G1/S genes, the average mRNA concentration (see above) was averaged over the 4 proB types above. Then, for all other cell types, a fold-enrichment relative to the pro-B group was calculated for each gene G as:

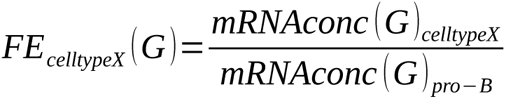

Fold-enrichments were then averaged over *CCND* genes, *CCNE* genes, *RB* genes, *E2F* genes *CDKN2* genes and *CDKN1A,C* genes to match the generic variables of the model. Finally, the model parameters that reflect the expression levels of those genes were renormalized based on the FE for each gene/cell type. Specifically, to model a cell type X, we updated model parameters as:

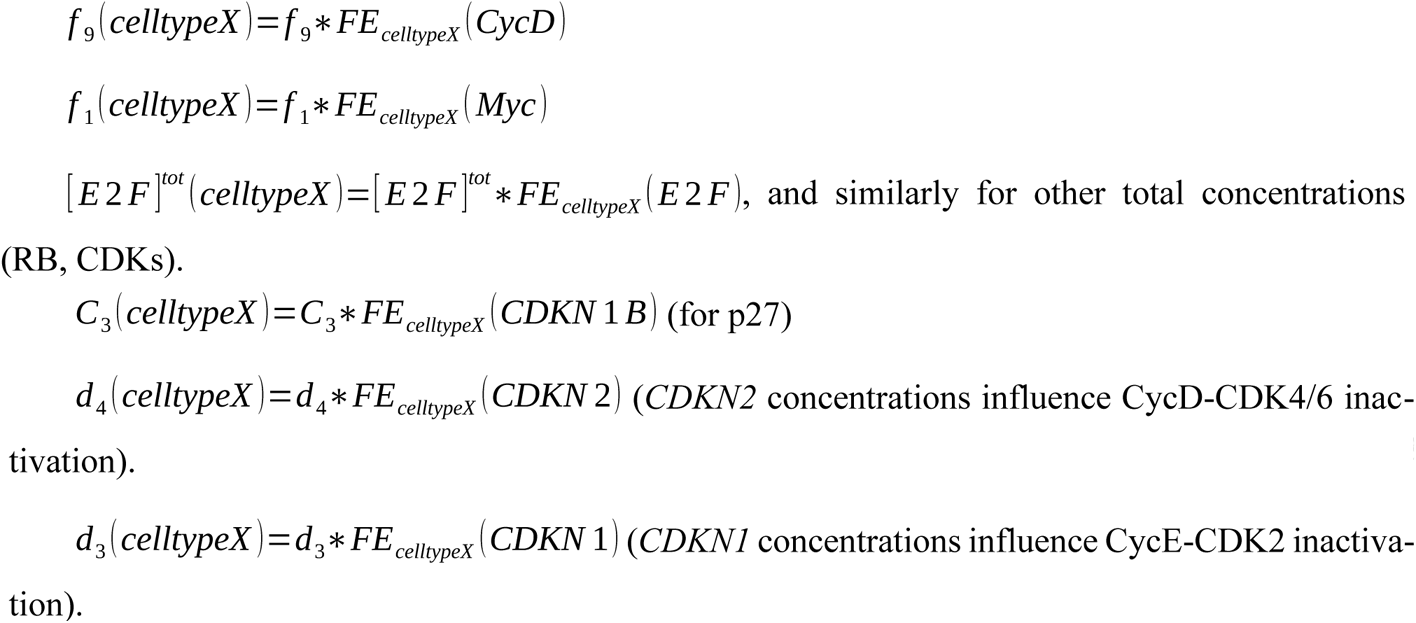

This data is reported in Supplementary Table T8 for all cell types.

### Model analysis

*Analytical resolution (partial) of steady state analysis and definition of the propensity to divide:* As detailed in the Supplementary Methods (Supplementary Figs. S1-S6), in the steady state the model ODEs reduce to a closed system of only 6 algebraic equations coupling 6 variables: [ *E* 2 *F* ]*^F^* /[ *E* 2 *F*_0_], the fraction of active E2F ; [*CDC* 25 *A* ]*^A^* /*Co*_1_, the concentration of active CDC25A relative to the threshold for CycE-CDK2 activation; _[_*Cyk _E_* _/ 2 ]_*^I^* /[*CDK* 2]*^tot^* , the fraction of catalytically inactive CycE-CDK2 complexes, relative to the total CDK2 concentration; _[_*Cyk _E_* _/2 ]_*^A^* /[*CDK* 2]*^tot^* , the fraction of catalytically active CycE-CDK2 complexes, relative to the total CDK2 concentration; _[_*Cyk _D_* _/4 ]_*^I^* /[*CDK* 4 ]*^tot^*, the fraction of inactive CycD-CDK4 complexes, relative to the total CDK4 concentration; and _[_*Cyk _D_* _/ 4 ]_*^A^* /[*CDK* 4 ]*^tot^* , the fraction of catalytically active CycD-CDK4 complexes, relative to the total CDK4 concentration. Those 6 steady state equations are parametrized by 16 meta-parameters, that combine one or several kinetic rates and other model parameters such that constant protein concentrations (e.g. CDKs, RB, E2F, Table 2). For the default parameters (pro-B cells, Table 2), the model was bistable (Supplementary Fig. S5-6), i.e. two stable steady state solutions co-existed and transitions between them were dynamically possible, following adequate stimuli. One state showed low activity (E2F mostly RB-bound, low CDK activity, low CDC25A concentration), and the other higher activity (E2F free, high CDK activity, high CDC25A concentration). This knowledge guided the numerical resolution of the equations presented below.

*Numerical resolution of the steady state equations:* steady state equations (Eqs.1-6, Supplementary Methods) were solved numerically using a fixed point methodology (Faires et Burden 2003). Briefly, a vector variable **x** is a fixed point of a function F if F(**x**)=**x**, which is the form of Eqs. 1-6. The fixed point algorithm choses arbitrary starting values for the variables, and iteratively modifies them using Eqs. 1-6, until the relative change between variables’ values between 2 iterations is less than a chosen tolerance (here 10^-6^). The algorithm was written in Matlab 2024b (Mathworks). Computations were run on a Lenovo 81wB laptop equipped with IntelCore I5 10210U CPU (1.6GHz) and 8Gb physical RAM. The algorithm was robust to changes in the starting values for the variables (Supplementary Table T5).

*Time-dependent simulations of the model:* to generate full time-dependent solutions of the model ODEs, we implemented dynamical numerical simulations with the Ode45 solver in Matlab R2024b (Mathsworks), that operates a Runge Kutta integration algorithm (Faires et Burden 2003). In presence of a bistable steady state, dynamical simulations were performed starting from two types of initial state: either “inactive”, where all proteins whose total concentration is known were assumed inactive and unbound, and all proteins whose concentration evolves in the model (CycE, CycD, CDC25A and MYC) were assumed to be absent; or “active”, where all the E2F pool was assumed RB-free, CycD, CycE, CDC25A and MYC concentration were fixed to 20-200 nM and most CDKs were cyclin-bound. The first and second sets of initial conditions converged respectively to the “low activity” and “high activity” steady states, we generated 1000 timecourses with random initial conditions and all converged to either of the steady states (Supplementary Table T6).

*Stability analysis:* we performed Lyapunov stability analysis using 100 randomized initial conditions picked in a +/-20% neighborhood of each steady state separately, and running the time-dependent simulations as described above. Convergence after 50.000 s is reported in Supplementary Table T7.

*Parameter space analysis:* a comprehensive numerical analysis of the effect of parameters on model outputs was performed using the 16 meta-parameters and steady state equations. 678978 parameter sets were randomly generated, using for each parameter 10^(-2+4*r)^ where r is a random number, uniformly distributed on [0,1] (with the exception of K_9_, that was randomly sampled on [0,1]). Hence, all meta-parameters but K_9_ varied between 0.01 and 100. The steady state was computed for the 6 variables defined above for each parameter set, using low activity initial conditions, generating a (16+6)*678978 data matrix. For each meta-parameter, the fraction of active E2F and parameter value were co-binned and the resulting 3D histogram was represented as colour-coded 2D plots (Supplementary Fig. S7).

*Sensitivity analysis:* we performed local sensitivity analysis (SA, (Hamby 1995)) to assess the effects of each meta-parameter individually on model output variables. we proceeded as follows: first, we computed the relative deviation in the steady state values of the biochemical variables ( *SS_new_*−*SS_default_* )/ *SS_default_* caused by a 1% change in one meta-parameter *kp*; then, we normalized this deviation to the relative change in this meta-parameter to its default value (Hamby 1995):

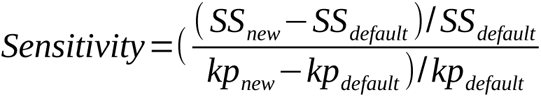

To prevent biases in sensitivity that would arise from widely different physiological ranges of distinct parameters, we converted most meta-parameters to log scale prior to calculation of the relative change, following what was done in parameter space analysis. This analysis was performed for 4 different total E2F concentrations.

*Global sensitivity analysis (GSA):* we used Pearson correlation coefficients (PCC, (Marino et al. 2008)) to capture linear trends between the 6 model outputs and the 16 meta-parameters. Specifically, pairwise Pearson correlations were computed between variables/meta-parameter, meta-parameter/meta-parameter and variable/variable pairs across the 678978 simulated samples (see above, parameter space analysis), and the correlation matrix was clustered using hierarchical clustering (clustergram function in Matlab 2024b, Pearson’s R coefficient and associated p-value, Supplementary Fig. S9).

### Model predicted propensity to divide

Due to the existence of multiple routes through G1/S, showing distinct requirements for E2F activation and CDK activation, we defined the propensity to divide for each cell type as the geometric mean of the total pool of active (RB-unbound) E2F and the total pool of active CDK (in the model, CDK2+CDK4 explicitly accounted for):

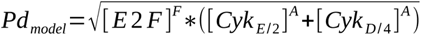

## Results

### Expression patterns of the 24 genes defining the core G1/S network recapitulate the diversity of immune cell types and of their proliferative properties

To explore how the expression of the core G1/S regulatory genes (Fig. 1; Table 1; Methods) varies across hematopoietic cell types and relates to their differential proliferation, we utilized the comprehensive human hematopoiesis atlas published by Zhang et al. (X. Zhang et al. 2024). As this scRNA-seq macro-dataset integrates datasets from several experiments, we minimized batch effects by focusing our analysis on the Human Cell Atlas (HCA) dataset (Hay et al. 2018) that encompasses scRNA-seq profiles from ∼100k cells from the bone marrow (BM) of 8 healthy adult donors.

Since cells that pass the G1/S checkpoint typically complete the cell cycle and divide (Donjerkovic et Scott 2000), we used the expression of essential S, G2 and M phase genes (Supplementary Table T2, (Macosko et al. 2015; Bertomeu et al. 2018)) to define a propensity to divide coefficient (P_d_, ranging from 0 to 1) for each of the 68 cell types (Methods). Throughout the manuscript, P_d_ will be our readout for proliferation. The inferred P_d_ was homogeneous across donors but strongly differed across the different cell types, highlighting erythroid-primed Megakaryocyte–Erythroid Progenitors (MEP-Eryth-1), late erythroid progenitors (ERPs) and erythroblasts, progenitor B (pro-B) cells and late monocyte-dendritic progenitors (MDPs) as the most proliferative hematopoietic cell types in the adult human bone marrow (Fig 2A, Supplementary Table T1). we partitioned hematopoietic cell types into 5 categories (mostly quiescent, low proliferation, mid, high and very high proliferation) using the Jenks algorithm. The P_d_ evolved similarly along most branches of the hematopoietic tree, with HSCs and early progenitors showing rather low P_d_ (∼0.05-0.2, quiescent/ low proliferation), later progenitors showing much higher P_d_ (0.25-1), and finally terminally differentiated cells showing very low P_d_ (0-0.1, Fig. 2B and 2C).

Since the G1/S checkpoint gates proliferation, we hypothesized that the regulation of the G1/S regulatory genes (Table 1) in consistent within cell types, forming lineage-specific expression patterns that reflect or even explain the distinct proliferative properties along the hematopoietic tree (Fig. 2).

To test this hypothesis, we aggregated cells from the same donor-cell type combination to form pseudobulk expression profiles, and performed unsupervised Principal Component Analysis (PCA) clustering based on the expression of the 24 core G1/S genes (Table 1). Because small gene sets can resolve only broad immune cell type classes (X. Chen, Chen, et Thomson 2022), we used the coarser (“Level 1”) annotations from Zhang et al. (Fig. 2A) encompassing 24 cell types (see Methods) to enable meaningful interpretation of the resulting clustering patterns.

The expression profiles of the G1/S gene set — despite its narrow mechanistic focus — clustered the 24 cell types (Fig. 3A, left) nearly as effectively as the 24 most highly variable genes (HVG) of the dataset (Fig. 3A, middle right), and way better than 24 randomly selected genes (Fig. 3A, right), as quantified by the clustering performance metrics Adjusted Silhouette width (ASw), Adjusted Rand Index (ARI) and (Normalized Mutual Information) NMI (Luecken et al. 2022)(Fig. 3B, right). In comparison, a list of 24 genes coding for receptors sensing bone marrow niche signals (Supplementary Table T3) (De Jong et al. 2024) performed only slightly better than the 24 G1/S genes in the task of clustering cell types across the 8 donors (Fig. 3A, middle left).

The variation in proliferative potential was best captured by the G1/S genes-based PCA, compared to HVG or hematopoietic receptors: the lead principal component (PC1) segregated well highly proliferative cell types (PC1 > 2.5) from less proliferative stem cells and early progenitors (0 < PC1 < 2.5), and terminally differentiated populations (PC1 < 0) (Fig. 3B, left). In contrast, PC2 mainly distinguished different branches of the hematopoietic tree, indicating the existence of distinct structures within the 24 G1/S network that would both affect hematopoietic proliferation and lineage identity.

Hence, together the expression pattern of the sole 24 core G1/S genes is sufficient to define hematopoietic cell types across all lineages, beyond inter-donor variability. This compact and interpretable gene set captures both lineage distinctions and proliferative dynamics of hematopoietic cells, supporting the critical role of the G1/S transition in coordinating division with differentiation. This observation also validates this list of core 24 G1/S genes for further mechanistic analysis of differential G1/S regulation across the hematopoietic tree.

### Three gene modules within the G1/S network are sequentially expressed along differentiation paths

The ability of G1/S genes to cluster diverse hematopoietic cell types effectively indicated the presence of distinct gene signatures within this core G1/S network which would be expressed in a lineage- and differentiation-stage specific way. This motivated us to interrogate whether co-expressed gene modules exist within the G1/S and display distinct expression patterns along the branches of the hematopoietic tree.

To identify co-expressed G1/S genes, we computed gene-gene Pearson correlations across the cell type*donor space. we used the fine annotation level that distinguishes 68 bone marrow hematopoietic cell types (Fig. 2) to increase the number of samples from which correlations are calculated, thereby improving the correlation estimates.

The resulting correlogram highlighted 3 gene groups with strong intra-group correlations and varying degrees of inter-group correlations (Fig. 4A, Supplementary Table T4). we refer to these 3 groups as functional “modules” within the G1/S network: a “Cyclin D module” encompassing *CCND1*, *CCND2*, *CDK6*, *MYC* and *E2F3*; a “Cyclin E module” consisting of *CCNE1*, *CCNE2*, *CDK2*, *CDKN3*, *CDC25A* (activator of CycE-Cdk2 complexes), *E2F1*, *E2F2*, *RBL1*, *CDK4* and *CDKN2C*; and a “Inhibitors” module comprising of *RB1*, *RBL2*, other *CDKN* genes (*CDKN1A*, *CDKN1B*, *CDKN1C*, *CDKN2A*, *CDKN2B*, *CDKN2D*) and, surprisingly, *CCND3*.

Intra-module correlations were strongest in the Cyclin E module, followed by the Inhibitors module and weaker, but still positive, in the Cyclin D module. The Cyclin D module showed consistent positive correlations with the Cyclin E module and negative correlations with the Inhibitors module. Interestingly, the Cyclin E module also correlated positively with the Inhibitors module at some developmental stages (Supplementary Fig. S11). E2F1 and E2F2 were co-expressed with the Cyclin E module, while E2F3 clustered into the Cyclin D module. we note that CDK4, which functionally should belong to the Cyclin D module, clustered within the Cyclin E module. However, *CDK4* expression showed strong correlations with all genes of the Cyclin D module as well, falling therefore in between these two modules.

we next assessed whether these 3 modules show distinct expression patterns across the hematopoietic tree. To examine how each cell type balances the expression between the 3 modules, we computed a “module score” for each cell type-donor sample (Methods) and projected these in barycentric coordinates (Fig. 4B). Cell types expressing predominantly Cyclin D, Cyclin E or Inhibitors module aligned with low, high/very high, and very low propensity to divide (Fig. 4B top).

The Cyclin D module dominated the expression of G1/S regulatory genes at the root of the hematopoietic tree (HSCs, MPPs), while the Cyclin E module dominated in strongly proliferating early to mid-stage progenitors. Interestingly, the gene most strongly correlated with the proliferative P_d_ score across all cell types was CDKN3 – a Cyclin E module gene and known CDK inhibitor (Supp. Fig. S10). In contrast, the Inhibitors module was strongly expressed in the quiescent, terminally differentiated cells across all lineages (Fig. 4B). Developmental transitions often coincided with shifts in module dominance, exemplified by the expression or shutdown of specific genes (Supplementary Fig. S11): for example, HSCs/CLPs express *CDKN2* family inhibitors at very low levels, while these inhibitors become strongly up-regulated in B-lymphoid progenitors (Pro-B). In parallel, a strong *CDK2* up-regulation and a down-regulation of *CDK4/6* drives the developmental trajectory of Pro-B cells toward the “Cyclin E – Inhibitors” wall (Fig. 4B, bottom triangle). Overlaying the modules’ activity to the hematopoietic landscape with single cell resolution (UMAP, Fig. 2B) confirmed these trends at the single cell level (Fig. 4C).

In conclusion, a sequential gene activation of three G1/S sub-modules marks hematopoietic differentiation: each branch of the hematopoietic tree begins with stem/stem-like cells with low propensity to divide that express predominantly the Cyclin D module genes; then these cells evolve to more proliferative progenitors that rely on the Cyclin E module, until Inhibitors dominate the expression of G1/S regulators and terminal differentiation is reached (Supplementary Fig. S11). The module sequence — Cyclin D, Cyclin E and finally Inhibitors — is consistent across hematopoietic lineages, while variations in the expression of the modules’ individual genes defined lineage identity (Supplementary Fig. S11).

To further understand the functional consequences on the G1/S transition of the uncovered hematopoietic G1/S gene expression dynamics, we sought to develop a mathematical model of the G1/S network that connects G1/S genes’ expression to E2F/CDK activation and the propensity to divide.

### A mathematical model of the G1/S transition across distinct hematopoietic cell types

To design a model able to capture proliferation properties of all hematopoietic cell types, we curated known interactions between the 24 G1/S genes’ products from the literature, and assembled them into a protein interaction network (Fig. 1). Our model includes one generic representative for E2F transcription factors (“E2F”), RB-like proteins (“RB”), D-type cyclins (“CycD”) and E-type cyclins (“CycE”). Moreover, the effects of CDK inhibitor proteins were incorporated through modulations of specific parameter (see Methods for details). This modeling framework can be readily extended to explicitly model the individual proteins within the generic representatives.

we described time-dependent changes in protein concentrations (via synthesis/degradation), assembly into complexes and activation/inactivation using a system of Ordinary Differential Equations (ODEs), based on mass action kinetics. we accounted for protein synthesis, degradation, activation, inactivation reactions, and binding/unbinding into complexes that can be activated/inactivated. Activation functions – such as the activation of CDC25A by CycE-CDK2 and the breakdown of the RBE2F complex by CDK activity – were modeled as Hill functions, to provide more flexibility in describing how outputs respond to inputs. The model’s outputs that are solved for are the concentrations of 15 interacting molecular species: MYC, free CycD ([*CyD* ]*^F^*), free CycE ([*CyE* ]*^F^*), RB-free E2F ( [ *E* 2 *F* ]*^F^*), active and inactive CDC25A ([*CDC* 25 *A* ]*^A^* and [*CDC* 25 *A* ]*^I^* ), unbound inactive CDKs ( [*CDK* 2]*^I^ ,* [*CDK* 4 ]*^I^ ,* [*CDK* 6 ]*^I^* ), active and inactive CycE-CDK2 complexes (_[_*Cyk _E_* _/ 2 ]_*^A^* and _[_*Cyk _E_* _/ 2 ]_*^I^* ), active and inactive CycD-CDK4 complexes (_[_*Cyk _D_* _/ 4 ]_*^A^* and _[_*Cyk _D_* _/ 4 ]_*^I^* ), and CycD-CDK6 complexes ( _[_*Cyk _D_* _/ 6 ]_*^A^* and _[_*Cyk _D_* _/ 6 ]_*^I^* ). Their inter-dependent dynamics are governed by a set of 38 model parameters: 26 kinetic rate parameters, 3 Hill exponents, 4 threshold concentrations and 5 total protein concentrations for RB, E2F and CDKs (Table 2). Full model equations are provided in Supplementary Methods, and default parameter values are listed in Table 2. Model construction and criteria for the choice of the default parameters are explained in the Methods section.

we first solved both the model’s dynamics (time-dependent variations) and its steady state computationally, using respectively Runge-Kutta integration and a robust fixed point algorithm (Supplementary Table T5, (Faires et Burden 2003)). with default parametrization (Table 2) and across random initial conditions, the model solutions consistently converged over time towards one of two discrete steady states: a “low activity” state (Fig. 5A), or “high activity” state (Fig. 5B). The low-activity state was characterized by RB-bound E2F, low CDK activity, low concentrations of CycE and CD - C25A, and inactive (“off”) Hill functions. In contrast, the high-activity state showed RB-free E2F, elevated CDK2 activity, over 10 fold higher CycE and CDC25A concentrations, and active (“on”) Hill functions. The convergence towards one or the other steady state depended on initial conditions: simulations beginning in a state with elevated CycD/CycE pools, some active Cyc-CDK complexes and/or active CDC25A resumed convergence towards fully active state, while simulations beginning with inactive Cyc-CDK complexes and without pre-existing CycD/E or CDC25A pools converged to the inactive steady state (Supplementary Table T6). Both steady states were dynamically stable in the Lyapunov sense, as perturbation of the steady state variables by +/20% of their value resulted in re-convergence to the same state. Thus, under default parameters, the G1/S transition exhibited robust bistability.

In summary, numerical simulations of the model using the default parameter set (Table 2) generated time courses for the concentration of G1/S proteins that evolved towards a low-activity state where the cell is not engaged in the G1/S transition or a high-activity state where E2F is activated and G1/S transcription is expected, committing the cell to division. Both states are stable to small perturbation (+/20 %), showing that the network structure protects cells from molecular noise. Transition from the low activity state to the high activity state therefore requires stronger perturbation, reflecting the binary decision a cell faces whether to remain in a non-dividing state, or commit to cell cycle entry. However, the default model parameters likely correspond to one cell type. Since different hematopoietic cell types express different levels of G1/S genes (Fig. 3-4, Supplementary Fig. S11), translating in into different model parameter regimes, we next sought to to gain mechanistic insight into how the 38 model parameters influence the model outputs. In this purpose, we performed a partial analytical resolution of the model’s steady state.

### Steady state analysis reveals 3 routes through the G1/S transition

E2F1-3 are together essential, at least in mouse embryo fibroblasts (wu et al. 2001); however, ectopic E2F activation is insufficient to induce S-phase in the absence of CycE (Duronio et al. 1996), and leukemia cell proliferation is reduced in vivo by CDK4/6 inhibition (Bride et al. 2021). Hence, both CDK activity and E2F activity (at least to a minimal level, and depending on the cellular context) are required to pass the G1/S transition, although both are intricately linked (Dong et al. 2018).

In an attempt to solve anaytically the steady state equations (Supplementary Methods Eqs. 1-6), we therefore first focused on analyzing E2F activation (Supplementary Methods, Eq. 1). In the steady state, the fraction of active E2F i.e. the ratio of the RB-free E2F concentration [ *E* 2 *F* ]*^F^*to the total concentration _[_ *E* 2 *F*_0 ]_ is given by:

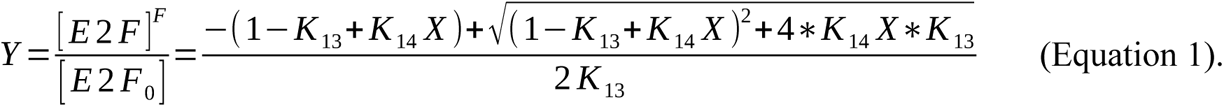

The meta-parameter 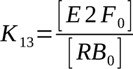 is the ratio of total E2F protein pool over the total RB protein pool. The meta-parameter 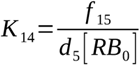 is the ratio of the maximal rate of RB-E2F dissociation (*f* _15_, corresponding to E2F activation, and that is achieved for complete RB phosphorylation), to the spontaneous RB-E2F binding rate *d*_5 [_ *RB*_0]_, that corresponds to E2F inactivation and is proportional to RB concentration. Because during the G1 and G1/S phases, RB phosphorylation is dependent on the concentration of catalytically active Cyc-CDK complexes, *K* _14_ is modulated by the fraction of phosphorylated RB-E2F complexes, which is assumed to be a Hill function of the total CDK activity (Methods, (Dorsey et al. 2018)):

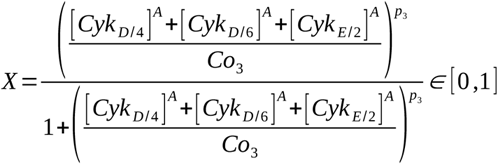. Here, the parameter *Co*_3_ is a threshold of active CDK concentration above which it efficiently catalyzes RB phosphorylation.

Plotting the active E2F fraction *Y* as a function of *K* _13_ and *K* _14_ *X* revealed three principal ways by which maximal E2F activation can be achieved (Supplementary Fig. S1): very large *K* _13_ (i.e., large excess of E2F vs RB), very large *K* _14_ (e.g. low RB concentration leading to unstable RB-E2F complexes); or *K* _14_ of order 1 or more, and maximal CDK activation *X* =1 . while the first two conditions might be achieved independently of any other model parameter, the third condition requires to fully solve the model for CDK activation (Supplementary Methods).

In total, we identified 6 distinct routes for E2F activation, and the triggering of the G1/S transi tion:

- **Route 1: E2F accumulation-driven G1/S transition.** when the total E2F concentration [ *E* 2 *F*_0_] greatly exceeds the total RB concentration [ *RB*_0_], we have *K* _13_ ≫ 1 and a large fraction of the E2F pool is active, without further requirement for an increase in CDK activity (Supplementary Fig. 1, bottom arrow). Thus, this route of E2F activation is independent of all other parameters.
- **Route 2: CycD-driven G1/S transition**. Even without strong E2F accumulation (relative to RB), a sufficient rise in the concentration of active CycD-CDK4 (_[_*Cyk* _]_*^A^*) or CycD-CDK6 complexes (_[_*Cyk* _]_*^A^*) increases the fraction of phosphorylated RB, which in turn activates the E2F pool, provided that, overall, *K* _14_ *X* >1. The higher *K* _14_ (e.g., low RB concentration), the less CDK activity ( *X* ) is required for E2F activation. Enhanced CycD-CDK4/6 activity, eliciting Route 2, can be achieved either *via* high CDK4/6 and CycD expression which directly increases the number of complexes (route 2a, Methods), or *via* sequestering by CycE-CDK2 of the common CDK inhibitor p27 which leads to CycD-CDK4/6 de-repression (route 2b). Route 2 does not require CycE-CDK2 activity, although CycE-CDK2 complexes may get activated downstream, as a consequence of E2F activation.
- **Route 3: CycE-driven G1/S transition.** In the more general case where CycD-CDK4/6 activity is insufficient on its own to fully activate E2F, CycE-CDK2 activity is required. Our model shows that an active CycE-CDK2 pool _[_*Cyk* _]_*^A^* can accumulate through a positive feedback loop in which E2F stimulates CycE synthesis (Supplementary Figs. S5, S6). This can occur through 3 sub-mechanisms: either with (route 3a) or without (route 3b) contribution from CDC25A, and/or or *via* sequestering of p27 by inactive CycD-CDK4/6 complexes, leading to CycE-CDK2 de-repression (route 3c).

we identified 16 meta-parameters, defined in Supplementary Methods and calculated in Table 2, whose value ranges determine which of the three G1/S routes defined above is triggered, as demonstrated mathematically in Supplementary Methods (Supplementary Figs. S1-6). Under the default parametrization, the meta-parameters *K* _13_ and *K* _14_were of order of magnitude 1, leading to a CDK-dependent E2F activation, excluding Route 1. Furthermore, the meta-parameters capturing CycD-dependent CycD-CDK4/6 binding and activation ( *K* _8_ and *K* _9_, Supplementary Methods Eq. 5) were much smaller than 1, making the residual CycD-CDK4/6 activity insufficient for E2F activation, also excluding Route 2. Instead, E2F activation under the default conditions required both the critical conditions 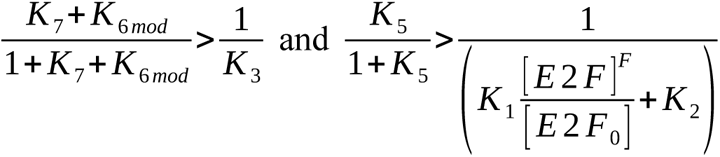 to be satisfied for higher E2F activity. Those conditions lead to a bistable, CycE-CDK2/CDC25A driven, steady state (route 3, Supplementary Figs. S5 and S6A).

E2F activation in the steady state was robust to variations in most individual meta-parameters over 4 orders of magnitude, as revealed by a comprehensive parameter space analysis (Supplementary Fig. S7). This robustness most likely stems from the strong redundancy built in the G1/S network, that offers multiple activation routes for both E2F and/or CDK activation (routes 1, 2a-b, 3a-b-c). Small values of one activatory meta-parameter which would block one activation route could be compensated by large values of other activatory meta-parameters, that enabled other activation routes. Thus, our analysis demonstrated that cells can preserve G1/S transition competence despite experiencing significant variations in molecular inputs. The analysis of correlations between meta-parameter values and model outputs (Supplementary Fig. S9) revealed that active CycD-CDK4/6 correlated positively with inactive CycE-CDK2 and vice-versa, indicating the relevance of the mechanism of competition for p27 across the whole meta-parameter space.

Since the meta-parameters depended on expression of the G1/S genes, hematopoietic transitions along the lineages are accompanied by changes in G1/S meta-parameters that may alter the dynamical modes through which different cell types pass the G1/S transition.

### G1/S gene expression patterns elicit distinct G1/S routes in distinct hematopoietic cell types

Having identified, in the mathematical analysis of our model, different routes to pass the G1/S transition that used either CycD and/or CycE and E2F or even E2F alone, we sought to predict whether those routes correspond to distinct hematopoietic cell types pass the transition. with default parameters, the model predicts low Cyclin D expression, low CycD-CDK4/6 activity (Fig. 5), little competition for p27 and a fully active CycE-CDK2/CDC25A feedback loop (route 3a, Supplementary Fig. S6A). To identify a cell type that matched those characteristics and may exemplify the default parameter, we looked for cell types with low *CCND* and *CDK4/6* expression, high *E2F* and *RB* expression, and relies on Cyclin E module for proliferation (on the CycE module-Inhibitors module wall on Fig. 4). we identified pro-B cells with such characteristics, and grouped pro-B-early-cycling, pro-B-cycling-1, 2 and transitional pro-B1 to reduce inter cell type variability in the pro-B pool (Methods). The mRNA concentrations of the 24 G1/S genes in other cell types were normalized to that of the pro-B group, and those normalized mRNA concentrations were used to rescale the corresponding model parameter(s) in other cell types proportionally, in order to model that cell type (Methods, Supplementary Table T8). For instance, the total CDK2 concentration in the model was rescaled to 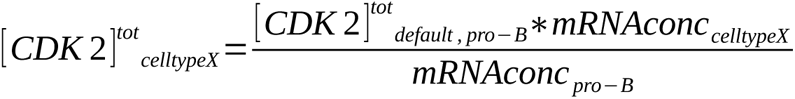

All 68 cell types were modeled this way (Supplementary Table T10), and we emphasized on ERP7, pre-DC-1, HSC/MPP (HSC2 + MPP1 + MPP2) and MEP-1 cells to illustrate our results. For each cell type, the meta-parameters were re-calculated from the rescaled model parameters, and the G1/S route was identified based on of those meta-parameter values, as analyzed in Supplementary Methods (Supplementary Table T9). All 5 highlighted cell types showed *K* _13_ *, K* _14_ >1 and a total CDK content larger than *Co*_3_, showing a priori potential for all G1/S routes.

while cycling pro-B cells were predicted to pass G1/S via route 3, ERP cells showed a fully active CycD-CDK4/6 with a strong contribution of the competition for p27 (route 2b), in addition to active CycE-CDK2. MEP-1 cells showed a E2F/RB ratio larger than 1, suggestive of route 1, yet also exhibited partial activation of both CycE-CDK2 and CycD-CDK4/6, alongside effects from the competition for p27 (route 1/3c). In contrast, pre-DC-1 and HSC/MPP cells showed a predicted monostable steady state, where the Cdc25-CycE feedback loop (Supplementary Fig. S5) was turned <OFF> (Supplementary Fig. S6D) and CycE-CDK2 activation was not rescued by the competition for p27

(unlike Supplementary Fig. S6F). In HSPCs, very low RB expression yielded a large 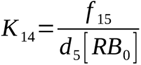 coefficient, ensuring minimal CDK activity (provided in those cells by CycD-CDK4/6 complexes, route 2a). In pre-DC-1 cells, high CycD expression was sufficient to provide CycD-CDK4/6 activation, despite a low E2F/RB ratio. Hence, our model predicts that evolution of G1/S genes expression across the hematopoietic tree affects the same G1/S molecular network encodes for different routes through the G1/S transition.

### The total amount of E2F is a key pivotal parameter that affects G1/S commitment across G1/S routes

Our analysis of the parameter space (Supplementary Fig. S7, *K* _13_) and global sensitivity analysis (Supplementary Fig. S9, highest correlation of active E2F with *K* _13_) indicated that the fraction of active E2F, our readout for the G1/S transition, depends most strongly of total E2F concentration more than on other parameter. This dependency holds across the entire parameter space and thus applies across all G1/S routes. In line with this result, we recently have shown that in yeast, accumulation of the E2F functional analog SBF to a threshold level during G1 was necessary to enable the G1/S transition (Dorsey et al. 2018; Goswami et al. 2025; Black et al. 2020). Motivated by these experimental findings, we next examined how varying total E2F concentration influences the steady state levels of the model variables. Keeping other meta-parameters fixed at default values, we found a sharp transition in Cyc-CDK activity: below the threshold[ *E* 2 *F*_0_ ]<0.85∗[ *RB*_0_ ], Cyc-CDK activity remained low, while above this threshold, a high activity was observed (Fig. 6A). Notably, the concentration of active CycE-CDK2 was much sensitive to E2F levels than to the concentration of CycD-CDK4/6 complexes.

At this tipping point, the model outputs – including the fractions of active E2F, active and inactive CycE-CDK2 complexes, and active CDC25A become hyper-sensitive to very small changes in two meta-parameters *K* _8_ and *K* _10_. A mere 1% change in either meta-parameter values yields about 25% change in output values (sensitivity analysis, Fig. 6B). As *K* _8_ and *K* _10_ represent respectively the activation rate of CycD-CDK4/6 complexes and the E2F-dependent CycE synthesis rate, this result indicate a dynamical instability that appears when sufficient E2F is accumulated: small fluctuations in CycE synthesis of CycD-CDK4/6 activation strongly affect the model’s activation. Sensitivity to CycD-CDK4/6 accumulation and activation was much lower, and restricted to E2F activation for _[_ *E* 2 *F*_0 ]_<0.85∗_[_ *RB*_0 ]_; and for _[_ *E* 2 *F*_0 ]_>0.85∗_[_ *RB*_0 ]_ the system’s steady state was almost insensitive to fluctuations in meta-parameters, emphasizing the robustness of the transition (Supplementary Fig. S8).

The pattern where E2F accumulation to a threshold 0.85 times the amount of RB for default parameters – was driving the model to a bistable state, hypersensitive to small fluctuations in CycE synthesis or CycD-CDK4/6 activation, was not encountered in all cells types. Modeled erythroid progenitors (such as ERP7) or erythroblasts for instance showed full E2F activation even for very low total E2F concentratios, due to the excessive CycD-CDK4/6 activity. Likewise, in HSCs, E2F activation did not require strong E2F accumulation, possibly due to very low RB levels; however, high E2F concentration was required to activate CycE-CDK2.

### B lymphoid opt for E2F-driven proliferation while MEP/ERP undergo CDK-driven proliferation

Our mathematical model identified the total pool of E2F as a pivotal parameter controlling the G1/S transition across routes, prompting us to examine whether *E2F* expression levels correlate with the propensity to divide across hematopoietic cell types. Because some cell types – such as late erythroid and B-cell progenitors – express preferentially the Cyclin E module (with *E2F1-2*), while others – such as HSC/MPPs and earliest myeloid progenitors (MultiLin-GMP) - favor the Cyclin D module (including *E2F3*) (Fig. 4 and Fig. S11), we considered the summed expression of *E2F1-3* (total *E2F*) in order to enable comparisons the data across cell types (Methods). we then plotted the propensity to divide P_d_ for each cell type against total *E2F* per cell (Fig. 6C, left).

The positive correlation between the propensity to divide P_d_ and total *E2F* transcript abundance per cell type was strong (r=0.79, p<0.01, Pearson test, Fig. 6C, left), with highest amounts in late ERP, Erythroblast, MkP-late, Pro-B-cycling-1 and Transitional-B-1 cells. However, significant dispersion appeared among cell types with mid-to-high *E2F* values – suggesting that additional factors beyond E2F transcript abundance contribute to P_d_ in this range.

Since differences in sequencing depth are mitigated by the use of pseudobulk samples, we hypothesized that this dispersion may essentially originate from large differences in the total RNA content across cell types, that would affect *E2F* RNA content. Therefore, we normalized the E2F counts to the total RNA content in each cell, defining a *E2F* concentration per cell (Methods). This normalization revealed that P_d_ scales with *E2F* concentration only in cell types with low P_d_ and *E2F* (r=0.41, p<0.01, Pearson test, Fig. 6C, right), Fig. 6C, right). In contrast, cell types with high P_d_ displayed either moderate *E2F* concentration (MEPs and ERPs) or very large *E2F* concentration (pro-B cells). These results suggest that while a minimum *E2F* concentration is required for proliferation, exceeding this threshold does not further enhance P_d_.

Our model predicted that, as an alternative to E2F driven G1/S commitment (route 1, and to a lesser extent route 3), cell could pass through G1/S by relying on excessive CDK activity (route 2, Supplementary Fig. S1). To test this, we plotted P_d_ for each donor/cell type pseudobulk sample against the total CDK concentration.

Overall, P_d_ correlated positively with CDK concentration, but certain cell types with mid/high P_d_ had low CDK concentrations (0-0.5), and instead had a E2F/RB ratio ≥1 (Fig. 7A). Thus, as for E2F, once a minimal CDK concentration is reached, the correlation between CDK concentration and P_d_ is weak. This trend was particularly evident in erythroid progenitors cell types (ERP1-8), which showed a rather high variability in Pd despite but quite similar total CDK concentrations, indicating that in these cells, other factors dominate the G1/S transition.

Since ERP cells express both CycD and CDK4/6 at high level (Supplementary Table T8, Supplementary Fig. S11), we hypothesized that in these cells, the limiting factor is rather the CycE-CDK2 activity, that requires CycE synthesis (under the control of E2F). To test this hypothesis, we compared the model-predicted CycE levels against the total *CCNE1-2* expression in individual ERP stages, and colored them according to their propensity to divide (Fig. 7B). The correlation between model-predicted CycE synthesis and CCNE1-2 expression in primary ERP cells was very high (r=0.86, p<0.01, Pearson test), and confirmed that higher CycE levels associate with higher Pd in ERPs. Hence, although ERPs express Cyclin D module genes at very high levels, the limiting factor for them to pass G1/S is CycE synthesis.

To integrate the contributions of both E2F- and CDK-driven mechanisms, we defined a unified model-based propensity to divide predicted by the model *P_d , model_* as the geometric mean of the total pool of active E2F and active CDK in the steady state (Methods). This definition captures all routes of G1/S activation. Solving the model across all 68 hematopoietic cell types yielded predicted proliferation propensities (Supplementary Table T10), which correlated well with the scRNA-seq data derived P_d_ (r=0.65, p<0.01, Pearson test), especially for low and mid proliferative cell types (Fig. 7C). The correlation was weaker in highly proliferative cell types, indicating that once minimal levels of E2F and CDK activity are attained, the propensity to divide is modulated by additional factors – either not included in our model, or the nature and quantitative effect of which is cell type-dependent.

In summary, the proliferation of hematopoietic cell types rely either on a sufficiently large E2F/RB ratio or elevated CDKs expression, consistent with the model’s prediction of E2F-driven (route 1 and 3) or CDK-driven (route 2) G1/S transition routes. This is captured by our model-derived propensity to divide *P_d , model_*. Once the inter-dependen E2F and CDK thresholds are met, the proliferative potential is further shaped by additional, potentially lineage-specific regulatory inputs.

### The routes through G1/S are elicited by G1-dependent concentration of distinct G1/S activators/ inhibitors across cell types

For most genes, mRNA production and degradation rates are coordinated with cell size to maintain mRNA concentration homeostasis – a phenomenon known as size-scaling (Berry et al. 2022). However, some G1/S regulators exhibit nonlinear scaling: while mRNAs of certain G1/S activators were accumulating faster relative to cell size (super-scaling), the opposite was true for G1/S repressors (sub-scaling, (Yuping Chen et al. 2020)). Differential scaling of G1/S activators/repressors with respect to cell growth in G1 was also reported at the protein level (Dorsey et al. 2018; Zatulovskiy et al. 2020; Litsios et al. 2022; Swaffer et al. 2021). This differential scaling effect shifts the balance between protein concentrations of G1/S activators and repressors, increasing the likelihood that cells proceed through G1/s as they become bigger in G1 phase.

Since UMI-based scRNA-seq data accurately reflect total transcript abundances per cell, and these correlate with cell size (Heumos et al. 2023; Kim et al. 2023), we utilized the here analyzed human bone marrow dataset to assess whether differential size-scaling effects could drive the G1/S transition also in human hematopoietic cells. For this pupose, we calculated the single-cell RNA concentrations of the 24 G1/S genes and binned the cells according to their total RNA, used here as a proxy for cell size. Then we computed for each gene the average RNA concentration within each bin. we focused on example cell types for which the model suggested distinct routes through G1/S: HSC/MPPs and pre-DC-1 (route 1), ERP-7 and ERP-8 (route 2b), and pro-B-1 cells (route 3).

Variability in RNA content was observed across cell types, with pro-B-1 cells having the lowest total RNA content, followed by HSC-2, MPP-2, pre-DC-1, and finally ERP-7/8 cells that showed the highest transcript abundance, but within cell types the total RNA content was more evenly distributed (Supplementary Fig. 12A). Thus, to align the different cell types by cell cycle phase, we binned cells separately within each type. we then examined the expression pattern of canonical S/G2/M markers - *MKI67* (Marker Of Proliferation Ki-67), *TOP2A* (topoisomerase 2A), *H4C3* (H4 Clustered Histone 3) and *TUBB* (tubulin). The concentration of these post-G1/S markers rose beyond the fourth bin, indicating that the first 3-4 bins were predominantly populated with cells in G1 and G1/S phases (Supplementary Fig. 12B).

In ERP-7 and ERP-8 cells – locating at the top of the Cyclin D/Cyclin E module wall (Fig. 4), and predicted to utilize CycD-CDK4/6 driven G1/S route 2b - *CCNE1* concentration super-scaled with cell size (Fig. 8, in red), indicating that late ERP stages might need to accumulate enough CycE to pass the transition. This observation is in line with the strong correlation between *CCNE* concentration across ERP developmental stages at the population (cell type)-level and their propensity to divide (Fig. 7B). In contrast, genes from the inhibitors module, such as *RB* and *CDKNs,* as well as *E2F* (data not shown) and *CDK4* concentrations remained mostly constant across the 4 first size bins, indicating size-independent expression in G1 and early S phase. Hence, although ERP cells express Cyclin D module genes at a much higher level than most other cell types, they might rely on CycE-CDK2 activation to pass the G1/S transition.

In pro-B-1 cells – which are predominant Cyclin E/Inhibitors module users, with minor contribution of cyclin D module (Fig. 4), and predicted G1/S route 3 users, *CDKN1B* mRNA concentration rapidly declined with increasing cell size (Fig. 8, blue), indicating a potential reduction of p27 protein levels during G1. This sub-scaling of *CDKN1B* suggests a progressive release of CDK repression when G1 cells grow. In parallel, *CDK4* gene expression super-scaled with size. Thus, although pro-B cells express Cyclin E module genes at a much higher level than most other cell types (Fig. 4, Supplementary Fig. S11), they might rely on CycD-CDK4 activation to pass the G1/S transition. In line with this observation, the pro-B type with the highest P_d_ – Pro-B-Early cycling – exhibited a larger contribution of Cyclin D module than other pro-B subtypes, while still maintaining a relatively high Cyclin E module expression (Fig. 4, Supplementary Fig. S11). Given the relatively low expression of *CCND* and *CDK4/6* genes at the population level, CycD-CDK4/6 activity might represent the limiting factor to pass G1/S in pro-B cells.

HSC2 cells, that displayed a low E2F/RB ratio at the pseudobulk level (Supplementary Table T8) showed *RB1* expression slightly decreasing in the first 4 bins, while MPP2 cells, that displayed a pseudobulk E2F/RB ratio around 1 (Fig. 7B), did not (Supplementary Fig. S12); likewise, pre-DC-1 cells, that showed low CDK expression (but mostly active due to high CycD as predicted by our model, Supplementary Table T9), had an unfavorable E2F/RB ratio at the peudobulk level (Supplementary Table T8) but showed an increase in E2F transcription in the first 3 bins (Supplementary Fig. S12). This data indicates that HSC2 and pre-DC-1, which need to increase their E2F/RB ratio, may use differential scaling of *E2F* and *RB1* gene expression (Yuping Chen et al. 2020) during G1 as a tool to pass the G1/S transition. However, in those cell types, the data was very variable, possibly to the small number of cells scored overall, and by the fact that a dominant fraction of those cells is not cycling, introducing a major component to mRNA expression variability that is not associated with the cell cycle.

In summary, analysis of concentration changes of G1/S regulators in relation to cells’ total mRNA content uncovers a subtle regulatory layer that may not be evident at the population level. G1 hematopoietic cells may rely on cell type–specific size scaling of G1/S regulators to coordinate their division rate with their developmental phase. while gene expression patterns at the population level reflect cell type’s overall ‘priming’ to proliferate (P_d_), and to pass the G1/S transition through a specific route, the actual commitment to divide at the single cell level may be triggered by scaling-induced shifts in the concentration of key activators and inhibitors, which can be captured in our model through cell size-dependent protein concentrations (e.g. Fig. 6A, for E2F).

## Discussion

Despite half a century of intense research on the cell cycle mechanism, since Leland Hartwell and co-workers identified the first cell cycle genetic mutants in budding yeast, it is still unknown how cells take the decision to divide at the molecular level.

### Summary of our findings

In this work, we combine data-driven and model-driven approaches to gain mechanistic insight into the G1/S transition across cell types, using the healthy hematopoietic system as a model. Our work bridges the gap between previous “generic” mathematical models of the G1/S transition, that do not include cell type differences and therefore are unable to probe the different routes through which different cell types pass G1/S (Bayleyegn et Govinder 2017; Hume, Dianov, et Ramadan 2020), and data-only based studies (e.g., (Riba et al. 2022; Sukys et Grima 2025)) that account for distinct cell cycle gene expression across cell types but lack the theoretical framework to functionally link gene expression patterns to proliferation. To the best of our knowledge, our model is the first mathematical model of cell cycle commitment parametrized with RNA seq data across donor-derived primary cells at distinct developmental stages.

we restricted our transcriptomics data analysis to the expression of 24 genes encoding for the core G1/S regulators. we show that this shortlist is sufficient to define/cluster hematopoietic cell types across all lineages and capture their propensity to divide beyond inter-donor variability and without accounting for additional cell type markers, such as lineage-specific receptors. Hence, despite of its narrow mechanistic focus, the set of 24 core G1/S regulators appears as an efficient discriminator of hematopoietic cell identity. Expression patterns highlighted three distinct modules within the G1/S network, encompassing respectively cyclin D-related genes, cyclin E related genes, and G1/S inhibitory genes. These modules are sequentially expressed along hematopoietic developmental trajectories: Cyclin D in slightly proliferative HSCs and early multipotent progenitors, Cyclin E in highly proliferative lineage-biased progenitors, and Inhibitors in quiescent terminally differentiated cells respectively, along all hematopoietic lineages. The concordant upregulation of the Inhibitors module with Cyclin E module in proliferative lymphoid and myeloid progenitor cells may seem counter-intuitive (Fig. 4B, Supplementary Fig. S11). Yet, 70 years ago in his seminal work (<THE Chemical Basis of Morphogenesis> 1952), Alan Turing already remarked that robust switch-like systems based on feedback regulation require both strong activation and strong inhibition. No cell type was found on the Cyclin D module – Inhibitors module axis (Fig. 4B), because since Cyclin D expression is not dependent on a (E2F-dependent) positive feedback loop, Cyclin D inhibition and activation are opposed to each other and do not work de concert, unlike activators and inhibitors of the Cyclin E module.

Integrating the transcriptomics data into a mathematical model constructed from known interactions between the core G1/S regulators reveals that modules’ expression profiles project the G1/S network into distinct functional modes, that correspond to a continuum of routes through the G1/S transition, that we classified in 6 broad categories. Moreover, cell growth during G1 leads to size-dependent changes in the mRNA concentration of key G1/S regulators, which could – in a cell type-specific manner trigger commitment to the given G1/S route for which the cell is primed.

Despite its simplicity, our model estimated the propensity to divide in 68 hematopoietic cell types relatively well. we speculate that this heterogeneity in how the G1/S transition is wired across hematopoietic cell types provides means for the cells to react differentially to the same chemical stimulation within the same bone marrow microenvionment, to adjust G1 duration to developmental needs. Predicting further how the different routes through G1/S precisely affect developmental states would require to couple our model with a model for cell differentiation, as in (Cho, Kuo, et Rockne 2022). In this work, Cho et al. model cell division at a coarse, phenomenological level that does only account for the proliferation rate and whether the division is symmetric or asymmetric.

### Limitations

while our model is based on protein concentrations, it was parametrized with data deduced from mRNA measurements. For one a given cell type - in contexts where protein translation and degradation rates are not modulated by external factors - it can be assumed that the number of RNA concentration is proportional to the protein concentration (Liu, Beyer, et Aebersold 2016). Hence, comparisons of mRNA concentrations between small and large cells of the same type that reflect different growth stages in G1, can be reasonably converted at the protein level in the model parametrization, although translation/degradation rates may evolve in G1 for some genes (Litsios et al. 2024). However, translation and degradation rates do vary between different hematopoietic cell types: for example, the rather slow-proliferating HSCs have typically low translation rates while the fast-proliferating erythroid progenitors are translationally very active (Signer et al. 2014). These differences could explain discrepancies between our model predicted P_d_ (based on protein concentrations) and the P_d_ across different cell types (based on mRNA counts). Moreover, recent proteomics studies indicate that for proteins that super-scale with size, enhanced translation rather than increased transcription – is driving their accumulation (Litsios et al. 2024). In contrast, protein sub-scaling in fast-growing hematopoietic cells is predominantly a result of dilution effects, rather than active degradation (Leduc et al. 2025). Thus, our observed transcriptional upregulation of sile G1/S activators in small to medium-sized cells may translate to even stronger changes in protein concentrations. Consistent with this, the scRNA-seq data-derived P_d_ was typically exceeding the model predicted P_d_ in fast proliferating erythroid-lineage cells, suggesting that we under-estimated the G1/S (activatory) proteins concentrations in ERP/Erythroblasts cells.

It is noteworthy that neither our transcriptomics analysis approach, that quantified S/G2/M mRNAs at the population (cell-type) level, nor our modeling approach that predicted E2F and Cyc-CDK activation at the single cell level but without a definite readout for cytokinesis, did quantify cell proliferation as such directly. Hence, moderately high levels of a given gene in the transcriptomics analysis might originate from either very high levels of the gene in a small fraction of the cells, or moderately high levels in most cells, or anything in between, making the mapping of cell types to model G1/S routes via renormalized gene expression debatable. However, both approaches converged in that if the model predicts for a given cell type a low P_d_, it means that it does not predict strong E2F/CDK activation when fed with typical gene expression values for this cell type, and therefore that this cell type is not "primed" for G1/S activation, indicating a low P_d_ in vivo. Despite model’s simplicity, predicted and data-inferred propensities to divide correlated rather well (Fig. 7C).

To define our model, we have focused on the 24 proteins that constitute the core of the G1/S network. This gene subset cluster hematopoietic cell types and their P_d_ in PCA as efficiently as hematopoietic receptors, or as the most variable genes, demonstrating 1) that there is a strong correlation between G1/S properties and cellular identity along the hematopoietic tree, and 2) that these 24 genes capture the essence of the G1/S transition in hematopoietic cells. Yet, we did not include other factors that may play a direct mechanistic role on the transition, such as other E2Fs and/or their dimerization partners (Fischer et al. 2022). Moreover, our transcriptomics data analysis revealed that *CCND3* and *E2F3* deviated from *CCND1-2* and *E2F1-2* in their expression patterns across cell types, emphasizing on their functional specificity (Saleban, Harris, et Poulter 2023). Our mathematical model does not include specific functions for the different *CCND* or *E2F* genes.

### Integration of our results to current knowledge and hypotheses raised by this work

The existence of multiple routes through G1/S has been known for a long time, although G1/S plasticity is often attributed to cancer cells (Bhat et al. 2024; Knudsen, witkiewicz, et Rubin 2024; Bruschini, Ciliberto, et Mancini 2020). Our work demonstrates that, in the healthy hematopoietic system, a variety of routes through G1/S are captured quantitatively by a unified model based on just 24 G1/S regulators.

Our current G1/S transition model does not account for genes’ specificities within the Cyclin D or E2F families. However, *CCND1-3* genes and *E2F1-3* genes had distinct expression patterns across cell types. *E2F3* – together with *CCND1-2*, *CDK6* and *MYC* – was up-regulated in stem and early progenitor cells, whereas *E2F1-2* dominated in later developmental stages (Supplementary Fig. 11), Interestingly, *CCND3* concentrations were highest in fast-proliferating pro-B stages as well as several differentiated lymphoid cells, such as plasma B cells, T and NK cells – which, unlike other mature blood cells, can re-enter the cell cycle massively upon contact with an antigen. Moreover CycD3 governs proliferation of lymphoid cells after they leave the bone marrow, such as of mature B cells in germinal centers of lymph nodes (Ramezani-Rad et al. 2020), and of late T cell progenitors in the thymus (thymocytes, (Sawai et al. 2012)). These findings suggest that emerging roles of specific E2F and Cyclin D genes (Saleban, Harris, et Poulter 2023) might need to be included into a refined version of the model.

The Skotheim laboratory found that increasing RB1 dosage increases cell size at the G1/S transition almost linearly (Zatulovskiy et al. 2020). In our model, although cell size is not explicitly represented, increasing RB1 concentration led to an almost linear increase in the concentration of the critical amount of E2F required to elicit the transition, when pro-B cells were simulated (as a ‘reference’, with default parameters). Hence, for this cell type, RB1 dosage – in fact, the E2F/RB1 ratio - is a critical parameter constraining the G1/S transition, paralleling the experimental finding (Zatulovskiy et al. 2020). Notably, those experiments were performed under CDK4/6 inhibition, placing the cells into a regulatory mode resembling the type 1/3 routes observed in pro-B cells. However, when we simulated erythroid progenitors (ERP-7/8) – where high concentration of CycD-CDK4/6 is found – with increasing amounts of RB1, the dynamics of the G1/S transition were much less modified . These results align with recent data showing that RB1 downregulation during G1 is only essential in cells born with low CycD-CDK4/6 activity, while cells born with high CDK activity could commit to division without the need to degrade their RB1 pool (S. Zhang et al. 2024). Hence, as captured by our model, elevated enough E2F/RB1 ratio or CDK activity can function as independent but not mutually exclusive routes through the G1/S checkpoint, characterizing distinct cell types.

Consistent with our predictions, HSCs rely on the Cyclin D module (i.e., *CCND1-2*, *CDK6*) to pass the G1/S transition (Knudsen, witkiewicz, et Rubin 2024), corresponding to route 2. From a kinetics perspective, route 2 does not require time-consuming synthesis and accumulation of E2F, cyclin E and CDC25A (both under E2F control), and therefore allows for a shorter G1 phase. This prediction is supported by (Mende et al. 2015) who demonstrated that short G1 duration is critical for preserving stemness in human HSCs and progenitor cells (HSPCs). Conversely, re-directing the G1/S dynamics to the Cyclin E module (route 3) lead to a reduction in stemness and loss of functional HSPCs. Likewise, our model predicts that MEP-1 progenitors (MKP precursors, (X. Zhang et al. 2024)) require more E2F accumulation to fully activate CycE-CDK2 than MEP-2 (ERP precursors, (X. Zhang et al. 2024)), predictive of a longer G1 phase. This prediction aligns with the results published by Lu et al., who showed that overexpressing both CDK4/cyclin D1 and CDK2/cyclin E in MEPs results in a shorter G1 phase and evolution of MEPs towards the erythroid lineage, rather than megakaryocytic lineage (Lu et al. 2018b). Conversely, pharmacological inhibition of CDK4/6 with palbociclib which shifts cells from route 2 to route 3 prolonged the G1 duration in MEPs and promoted their differentiation towards megakaryocytes. These findings have implications for stem cell and regenerative medicine research, demonstrating that modulation of the G1/S network could maintain stemness or direct lineage commitment in therapeutic contexts.

Leukemic cells can derive from HSC or multipotent progenitors, or from lineage committed progenitors (X. Zhang et al. 2024; George et al. 2016). Leukemic cells tend to overexpress *CDK6* and *CCND2*, both belonging to the Cyclin D module, in acute myeloid leukemia (AML) and B-acute lymphoid leukemia (B-ALL) (S. Zhang et al. 2024). Hence, CDK4/6 inhibitors - such as palbociclib, ribociclib, and abemaciclib – that are FDA-approved anti-cancer drugs used in the clinics to treat breast cancer, are currently tested in trials for leukemia (Bride et al. 2021). In the Zhang et al. hematopoietic atlas (X. Zhang et al. 2024), AML leukemia stem cells aligned with early progenitors such that MPP-2, MPP–MEP, LMPP-1, MDP-2 or MultiLin-GMP-2, all of which show high *CDK6* and *CCND2* expression (Supplementary Fig. S11), and locate on the Cyclin D – Cyclin E module axis (Fig. 4B). when we model these cell types, they pass G1/S through route 2, characterized by constitutive E2F activity driven by elevated CycD-CDK activity. In this context, further CycE-CDK2 activation is possible upon accumulation of E2F, that increases CycE synthesis as in ERPs (Fig. 7B). In the light of our results, leukemic cells may undergo partial reversal of their development, as they turn from a progenitor state (Cyclin E module user) into a more HSPC profile (Cyclin D module user). Thus, according to our model, CDK4/6 inhibitors may impair leukemic cells while providing a competitive growth advantage to healthy cells that operate on the Cyclin E module - Inhibitors module axis (Fig. 4B), including pro-B, pre-DC and Mono-2 cells. This differential sensitivity to the drug may underlie the observed benefit of inhibiting CDK4/6 in T-ALL and B-ALL mouse models (Bride et al. 2021).

Moreover, our model’s steady-state – particularly the fraction of activated E2F and Cyc–CDK complexes – is poorly sensitive to changes in most meta-parameter value, indicating that the G1/S network is very robust to perturbation, such as fluctuations in protein expression or signaling. This robustness is conferred by the E2F-CycE-CDC25A positive feedback loops, which buffer against molecular noise. In contrast, the Cyclin D module responds linearly to perturbation. As a result, in the absence of additional mutations such as *CDKN2A* or *RB1*, lymphoblastic leukemia cells (Cyclin D module users) are predicted to be more sensitive than their healthy lymphoid progenitors (Cyclin E module users) to perturbations in the environment. This vulnerability could explain the strong selective pressure for mutations in *CDKN2A* and *RB1*—two of the most frequently inactivated cell cycle genes in leukemia (wang et Zhang 2025). *CDKN2A* loss of function dopes the Cyclin D route for G1/S activation (route 2), while *RB1* loss of function elicits the routes 1 and 3 by increasing the model’s meta-parameters K_13_ and K_14_. Hence, simultaneous *CDKN2A* and *RB1* mutation provides leukemic cells with plasticity for G1/S commitment, as all routes through G1/S get promoted, and the theoretical benefit of inhibiting CDK4/6 may be circumvent in leukemia.

Indeed, in other cancers the promising therapeutic effects of CDK4/6 inhibition are counterbalanced by the development of resistance to these drugs, that limits long-term efficacy (Kim et al. 2023). MCF-7 breast cancer cells treated with Palbociclib resumed growth following a 2-step process where RB was first degraded, followed by c-MYC-mediated E2F activation. The rapid degradation or RB upon CDK4/6 inhibition may be explained by the removal of CDK4-mediated hypophosphorylation of RB, that protects it from degradation (S. Zhang et al. 2024). In parallel, c-MYC activation promotes CDC25A synthesis and enables cells to bypass CDK4/6 inhibition through activation of the CycE-CDK2/CDC25A positive feedback (route 3). Hence, blocking route 2 via CDK4/6 inhibition is counterbalanced by c-MYC dependent increased activation of route 3. Our model explains therefore the strong correlation between patients’ c-MYC levels and their (poor) survival outcomes following CDK4/6 inhibition. In (Kim et al. 2023), it was suggested that restoring RB levels via proteasome inhibition could prevent relapse following CDK4/6 inhibition. Our model supports this hypothesis, predicting that increasing RB1 levels alters routes 1 and 3. Consistently, single breast tumor cells with elevated levels of E2F1, RB1 and CDK2 - key components of the Cyclin E module, priming to use routes 1 and 3 - tend to show stronger resistance to Pablociclib (Zikry et al. 2024), emphasizing the plasticity of the G1/S network.

CDK2 inhibition has also been tested in vitro against cancer cells, and is currently investigated in clinical trials against leukemia (House et al. 2025; Arora et al. 2023). However, resistance mechanisms have also emerged. Treatment of breast cancer cell line MCF10A with the PF3600 - selective for CDK2 at low concentrations (25–100 nM) and also targeting CDK4/6 at higher doses (500 nM) - led to a rapid decline in CDK2 activity. However, the initial drop was transient: within 1–2 hours, a compensatory CycD–CDK4/6 activation (route 2) recovered the CDK2 activity, allowing cells to complete the cell cycle. Our model provides a mechanistic explanation for this adaptative response: CDK2 inhibition is equivalent to increase the CycE-CDK2 de-activation rate d_3_, reducing K_6_ and K_7_ and therefore moving the blue curve downwards on Supplementary Fig. S5. Additional accumulation of E2F may compensate for this effect, by re-increasing K_6_ and K_7_, unless partial E2F activation is further compromised by additional inhibition of CDK4/6, as achieved with concomitant Palbociclib treatment in (Arora et al. 2023). Measuring E2F total expression and activation following CDK2 inhibition in MCF10A cells would validate or invalidate this hypothesis.

In summary, our model provides a quantitative analysis of the different routes that cells can take through the G1/S transition. Based on our analysis of hematopoietic cells, we propose that differential wiring of the G1/S network across cell types provides means for the cells to react differentially to the same biochemical environmental stimulation – being it growth signals from the bone marrow microenvironment, or pharmacological interventions -, and enables cells to adjust their G1 duration according to their developmental stage. we find that, although 6 different routes to pass G1/S can be distinguished, those are largely overlapping and provide a continuum of mechanisms. This work demonstrates that the commitment to division must be tackled at the systems level, and that one may never find a single mechanism of G1/S commitment. Our model is able to correctly recapitulate the wealth of propensities to divide across the hematopoietic tree. we envision that our model could be extended to other cell types - including cancer cells sampled from patients. By quantifying the RNA, or ideally the absolute protein concentrations, of the 24 core G1/S regulators, patient-specific renormalization of model parameters could enable tailored predictions of the efficacy of CDK or E2F inhibition strategies.

## Acknowledgments & funding statement

we thank the Bioinformatics Centre at University of Eastern Finland for access to computing facilities. This work was funded by the Academy of Finland (grant #350887 to ST) and the Sigrid Jusélius Foundation (grant #220196 to ST). A.H. conducted this project with personal funding from the Instrumentarium Science Foundation (grant #220006) and Finnish Cultural Foundation (grant #00240437) independent of the author’s primary research work. The funders had no role in study design, data collection and analysis, decision to publish, or preparation of the manuscript.

## Authors contributions

Conceptualization, S.T., A.H.; methodology, S.T., A.H, A.A.; software, A.A., A.H.; formal analysis, A.A., A.H., S.T.; writing—original draft preparation, S.T., A.H., A.A.; writing—review and editing, all authors; funding acquisition, S.T., A.H.. All authors have read and agreed to the published version of the manuscript.

## Abbreviations

CycD,E: cyclin D, E proteins;

## References

Ahmadi, Seyed Esmaeil, Samira Rahimi, Bahman Zarandi, Rouzbeh Chegeni, et Majid Safa. 2021. «MYC: A Multipurpose Oncogene with Prognostic and Therapeutic Implications in Blood Malignancies». Journal of Hematology & Oncology 14 (1): 121. 10.1186/s13045-021-01111-4.

Aibar, Sara, Carmen Bravo González-Blas, Thomas Moerman, Vân Anh Huynh-Thu, Hana Imrichova, Gert Hulselmans, Florian Rambow, et al. 2017. «SCENIC: Single-Cell Regulatory Network Inference and Clustering». Nature Methods 14 (11): 1083–86. 10.1038/nmeth.4463.

Arora, Mansi, Justin Moser, Timothy E. Hoffman, Lotte P. watts, Mingwei Min, Monica Musteanu, Yao Rong, et al. 2023. «Rapid Adaptation to CDK2 Inhibition Exposes Intrinsic Cell-Cycle Plasticity». Cell 186 (12): 2628–2643.e21. 10.1016/j.cell.2023.05.013.

Attwooll, Claire, Eros Lazzerini Denchi, et Kristian Helin. 2004. «The E2F Family: Specific Functions and Overlapping Interests». The EMBO Journal 23 (24): 4709-16. 10.1038/sj.emboj.7600481.

Bayleyegn, Y. N., et K. S. Govinder. 2017. «Mathematical Description of the Interactions of CycE/Cdk2, Cdc25A, and P27^Kip1^ in a Core Cancer Subnetwork». Mathematical Methods in the Applied Sciences 40 (8): 2961–79. 10.1002/mma.4213.

Berry, Scott, Micha Müller, Arpan Rai, et Lucas Pelkmans. 2022. «Feedback from Nuclear RNA on Transcription Promotes Robust RNA Concentration Homeostasis in Human Cells». Cell Systems ,13 (6): 454-470.e15. 10.1016/j.cels.2022.04.005.

Bertoli, Cosetta, Jan M. Skotheim, et Robertus A. M. de Bruin. 2013. «Control of Cell Cycle Transcription during G1 and S Phases.» Nature Reviews. Molecular Cell Biology 14 (8): 518-28. 10.1038/nrm3629.

Bertomeu, Thierry, Jasmin Coulombe-Huntington, Andrew Chatr-Aryamontri, Karine G. Bourdages, Etienne Coyaud, Brian Raught, Yu Xia, et Mike Tyers. 2018. «A High-Resolution Genome-wide CRISPR/Cas9 Viability Screen Reveals Structural Features and Contextual Diversity of the Human Cell-Essential Proteome». Molecular and Cellular Biology 38 (1): e00302-17. 10.1128/MCB.00302-17.

Bhat, Gh Rasool, Itty Sethi, Hana Q. Sadida, Bilal Rah, Rashid Mir, Naseh Algehainy, Ibrahim Altedlawi Albalawi, et al. 2024. «Cancer Cell Plasticity: From Cellular, Molecular, and Genetic Mechanisms to Tumor Heterogeneity and Drug Resistance». Cancer Metastasis Reviews 43 (1): 197–228. 10.1007/s10555-024-10172-z.

Black, Labe, Sylvain Tollis, Guo Fu, Jean-Bernard Fiche, Savanna Dorsey, Jing Cheng, Ghada Ghazal, et al. 2020. «G1/S Transcription Factors Assemble in Increasing Numbers of Discrete Clusters through G1 Phase». The Journal of Cell Biology 219 (9): e202003041. 10.1083/jcb.202003041.

Bride, Karen L., Hai Hu, Anastasia Tikhonova, Tori J. Fuller, Tiffaney L. Vincent, Rawan Shraim, Marilyn M. Li, et al. 2021. «Rational drug combinations with CDK4/6 inhibitors in acute lymphoblastic leukemia». Haematologica 107 (8): 1746–57. 10.3324/haematol.2021.279410.

Bruschini, Sara, Gennaro Ciliberto, et Rita Mancini. 2020. «The Emerging Role of Cancer Cell Plasticity and Cell-Cycle Quiescence in Immune Escape». Cell Death & Disease 11 (6): 471. 10.1038/s41419-020-2669-8.

Burda, Isabella, et Adrienne H. K. Roeder. 2022. «Stepping on the Molecular Brake: Slowing down Proliferation to Allow Differentiation». Developmental Cell 57 (5): 561-63. 10.1016/j.devcel.2022.02.014.

Butler, Andrew, Paul Hoffman, Peter Smibert, Efthymia Papalexi, et Rahul Satija. 2018. «Integrating Single-Cell Transcriptomic Data across Different Conditions, Technologies, and Species». Nature Biotechnology 36 (5): 411-20. 10.1038/nbt.4096.

Calegari, Federico, et wieland B. Huttner. 2003. «An Inhibition of Cyclin-Dependent Kinases That Lengthens, but Does Not Arrest, Neuroepithelial Cell Cycle Induces Premature Neurogenesis». Journal of Cell Science 116 (Pt 24): 4947-55. 10.1242/jcs.00825.

Carlberg, Carsten, et Eunike Velleuer. 2022. «Cells and Tissues of the Immune System». In Molecular Immunology, par Carsten Carlberg et Eunike Velleuer, 1-18. Cham: Springer International Publishing. 10.1007/978-3-031-04025-2_1.

Chen, Xiaoqiao, Sisi Chen, et Matt Thomson. 2022. «Minimal Gene Set Discovery in Single-Cell mRNA-Seq Datasets with ActiveSVM». Nature Computational Science 2 (6): 387-98. 10.1038/s43588-022-00263-8.

Chen, Yunshun, Lizhong Chen, Aaron T. L. Lun, Pedro L. Baldoni, et Gordon K. Smyth. 2025. «edgeR v4: Powerful Differential Analysis of Sequencing Data with Expanded Functionality and Improved Support for Small Counts and Larger Datasets». Nucleic Acids Research 53 (2): gkaf018. 10.1093/nar/gkaf018.

Chen, Yuping, Gang Zhao, Jakub Zahumensky, Sangeet Honey, et Bruce Futcher. 2020. «Differential Scaling of Gene Expression with Cell Size May Explain Size Control in Budding Yeast». Molecular Cell 78 (2): 359-370.e6. 10.1016/j.molcel.2020.03.012.

Cheung, Tom H., et Thomas A. Rando. 2013. «Molecular Regulation of Stem Cell Quiescence». Nature Reviews. Molecular Cell Biology 14 (6): 329-40. 10.1038/nrm3591.

Chinnam, Meenalakshmi, et David w. Goodrich. 2011. «RB1, Development, and Cancer». Current Topics in Developmental Biology 94:129-69. 10.1016/B978-0-12-380916-2.00005-X.

Cho, Heyrim, Ya-Huei Kuo, et Russell C. Rockne. 2022. «Comparison of Cell State Models Derived from Single-Cell RNA Sequencing Data: Graph versus Multi-Dimensional Space». Mathematical Biosciences and Engineering: MBE 19 (8): 8505-36. 10.3934/mbe.2022395.

Dalton, Stephen. 2015. «Linking the Cell Cycle to Cell Fate Decisions». Trends in Cell Biology 25 (10): 592–600. 10.1016/j.tcb.2015.07.007.

De Jong, Madelon M. E., Lanpeng Chen, Marc H. G. P. Raaijmakers, et Tom Cupedo. 2024. «Bone Marrow Inflammation in Haematological Malignancies». Nature Reviews Immunology 24 (8): 543-58. 10.1038/s41577-024-01003-x.

Dong, Peng, Carolyn Zhang, Bao-Tran Parker, Lingchong You, et Bernard Mathey-Prevot. 2018. «Cyclin D/CDK4/6 Activity Controls G1 Length in Mammalian Cells». PloS One 13 (1): e0185637. 10.1371/journal.pone.0185637.

Donjerkovic, D., et D. w. Scott. 2000. «Regulation of the G1 Phase of the Mammalian Cell Cycle». Cell Research 10 (1): 1–16. 10.1038/sj.cr.7290031.

Dorsey, Savanna, Sylvain Tollis, Jing Cheng, Labe Black, Stephen Notley, Mike Tyers, et Catherine A. Royer. 2018. «G1/S Transcription Factor Copy Number Is a Growth-Dependent Determinant of Cell Cycle Commitment in Yeast». Cell Systems 6 (5): 539-554.e11. 10.1016/j.cels.2018.04.012.

Duronio, R J, A Brook, N Dyson, et P H O’Farrell. 1996. «E2F-Induced S Phase Requires Cyclin E.» Genes & Development 10 (19): 2505-13. 10.1101/gad.10.19.2505.

Dyer, Sarah C., Olanrewaju Austine-Orimoloye, Andrey G. Azov, Matthieu Barba, If Barnes, Vianey Paola Barrera-Enriquez, Arne Becker, et al. 2025. «Ensembl 2025». Nucleic Acids Research 53 (D1): D948–57. 10.1093/nar/gkae1071.

Faires, John Douglas, et Richard L. Burden. 2003. Numerical Methods. 3. ed. Pacific Grove, CA: Thomson [u.a.].

Fischer, Martin, Amy E. Schade, Timothy B. Branigan, Gerd A. Müller, et James A. DeCaprio. 2022. «Coordinating Gene Expression during the Cell Cycle». Trends in Biochemical Sciences 47 (12): 1009-22. 10.1016/j.tibs.2022.06.007.

Galaktionov, K., X. Chen, et D. Beach. 1996. «Cdc25 Cell-Cycle Phosphatase as a Target of c-Myc». Nature 382 (6591): 511–17. 10.1038/382511a0.

Galloway, Kate E. 2024. «Changes in Cell-Cycle Rate Drive Diverging Cell Fates». Nature Reviews Genetics 25 (6): 379–379. 10.1038/s41576-024-00714-0.

George, Joshy, Asli Uyar, Kira Young, Lauren Kuffler, Kaiden waldron-Francis, Eladio Marquez, Duygu Ucar, et Jennifer J. Trowbridge. 2016. «Leukaemia Cell of Origin Identified by Chromatin Landscape of Bulk Tumour Cells». Nature Communications 7 (1): 12166. 10.1038/ncomms12166.

Gombos, Magdolna, Cécile Raynaud, Yuji Nomoto, Eszter Molnár, Rim Brik-Chaouche, Hirotomo Takatsuka, Ahmad Zaki, et al. 2023. «The Canonical E2Fs Together with RETINOBLASTOMA-RELATED Are Required to Establish Quiescence during Plant Development». Communications Biology 6 (1): 903. 10.1038/s42003-023-05259-2.

Goswami, Pooja, Abhishek Ghimire, Carleton Coffin, Jing Cheng, Jasmin Coulombe-Huntington, Ghada Ghazal, Yogitha Thattikota, et al. 2025. «Swi4-Dependent SwI4 Transcription Couples Cell Size to Cell Cycle Commitment». iScience 28 (3): 112027. 10.1016/j.isci.2025.112027.

Grinenko, Tatyana, Anne Eugster, Lars Thielecke, Beáta Ramasz, Anja Krüger, Sevina Dietz, Ingmar Glauche, et al. 2018. «Hematopoietic Stem Cells Can Differentiate into Restricted Myeloid Progenitors before Cell Division in Mice». Nature Communications 9 (1): 1898. 10.1038/s41467-018-04188-7.

Gu, Zuguang. 2022. «Complex Heatmap Visualization». iMeta 1 (3): e43. 10.1002/imt2.43.

Hamby, D. M. 1995. «A Comparison of Sensitivity Analysis Techniques». Health Physics 68 (2): 195-204. 10.1097/00004032-199502000-00005.

Hay, Stuart B., Kyle Ferchen, Kashish Chetal, H. Leighton Grimes, et Nathan Salomonis. 2018. «The Human Cell Atlas Bone Marrow Single-Cell Interactive web Portal». Experimental Hematology 68 (décembre):51-61. 10.1016/j.exphem.2018.09.004.

Hendler, Adi, Edgar M. Medina, Nicolas E. Buchler, Robertus A. M. de Bruin, et Amir Aharoni. 2018. «The Evolution of a G1/S Transcriptional Network in Yeasts.» Current Genetics 64 (1): 81-86. 10.1007/s00294-017-0726-3.

Henley, Shauna A., et Frederick A. Dick. 2012. «The Retinoblastoma Family of Proteins and Their Regulatory Functions in the Mammalian Cell Division Cycle». Cell Division 7 (1): 10. 10.1186/1747-1028-7-10.

Heumos, Lukas, Anna C. Schaar, Christopher Lance, Anastasia Litinetskaya, Felix Drost, Luke Zappia, Malte D. Lücken, et al. 2023. «Best Practices for Single-Cell Analysis across Modalities». Nature Reviews. Genetics 24 (8): 550–72. 10.1038/s41576-023-00586-w.

Hoffmann, I., G. Draetta, et E. Karsenti. 1994. «Activation of the Phosphatase Activity of Human cdc25A by a Cdk2-Cyclin E Dependent Phosphorylation at the G1/S Transition». The EMBO Journal 13 (18): 4302–10. 10.1002/j.1460-2075.1994.tb06750.x.

House, Isabelle, Mari Valore-Caplan, Elijah Maris, et Gerald S. Falchook. 2025. «Cyclin Dependent Kinase 2 (CDK2) Inhibitors in Oncology Clinical Trials: A Review». Journal of Immunotherapy and Precision Oncology 8 (1): 47-54. 10.36401/JIPO-24-22.

Hume, Samuel, Grigory L. Dianov, et Kristijan Ramadan. 2020. «A Unified Model for the G1/S Cell Cycle Transition». Nucleic Acids Research 48 (22): 12483-501. 10.1093/nar/gkaa1002.

Jeffrey, P. D., L. Tong, et N. P. Pavletich. 2000. «Structural Basis of Inhibition of CDK-Cyclin Complexes by INK4 Inhibitors». Genes & Development 14 (24): 3115–25. 10.1101/gad.851100.

Kim, Sungsoo, Jessica Armand, Anton Safonov, Mimi Zhang, Rajesh K. Soni, Gary Schwartz, Julia E. McGuinness, et al. 2023. «Sequential Activation of E2F via Rb Degradation and C-Myc Drives Resistance to CDK4/6 Inhibitors in Breast Cancer». Cell Reports 42 (11): 113198. 10.1016/j.celrep.2023.113198.

Knudsen, Erik S., Agnieszka K. witkiewicz, et Seth M. Rubin. 2024. «Cancer Takes Many Paths through G1/S». Trends in Cell Biology 34 (8): 636-45. 10.1016/j.tcb.2023.10.007.

Konagaya, Yumi, David Rosenthal, Nalin Ratnayeke, Yilin Fan, et Tobias Meyer. 2024. «An Intermediate RbE2F Activity State Safeguards Proliferation Commitment». Nature 631 (8020): 424-31. 10.1038/s41586-024-07554-2.

Kossatz, Uta, Kai Breuhahn, Benita wolf, Matthias Hardtke-wolenski, Ludwig wilkens, Doris Steinemann, Stephan Singer, et al. 2010. «The Cyclin E Regulator Cullin 3 Prevents Mouse Hepatic Progenitor Cells from Becoming Tumor-Initiating Cells». The Journal of Clinical Investigation 120 (11): 3820–33. 10.1172/JCI41959.

Kueh, Hao Yuan, Ameya Champhekar, Stephen L. Nutt, Michael B. Elowitz, et Ellen V. Rothenberg. 2013. «Positive Feedback between PU.1 and the Cell Cycle Controls Myeloid Differentiation». Science (New York, N.Y.) 341 (6146): 670-73. 10.1126/science.1240831.

Leduc, Andrew, Shanshan Zheng, Purvi Saxena, et Nikolai Slavov. 2025. «Impact of Protein Degradation and Cell Growth on Mammalian Proteomes». bioRxiv: The Preprint Server for Biology, avril, 2025.02.10.637566. 10.1101/2025.02.10.637566.

Litsios, Athanasios, Pooja Goswami, Hanna M. Terpstra, Carleton Coffin, Luc-Alban Vuillemenot, Mattia Rovetta, Ghada Ghazal, et al. 2022. «The Timing of Start Is Determined Primarily by Increased Synthesis of the Cln3 Activator Rather than Dilution of the whi5 Inhibitor». Molecular Biology of the Cell 33 (5): rp2. 10.1091/mbc.E21-07-0349.

Litsios, Athanasios, Benjamin T. Grys, Oren Z. Kraus, Helena Friesen, Catherine Ross, Myra Paz David Masinas, Duncan T. Forster, et al. 2024. «Proteome-Scale Movements and Compartment Connectivity during the Eukaryotic Cell Cycle». Cell 187 (6): 1490-1507.e21. 10.1016/j.cell.2024.02.014.

Liu, Yansheng, Andreas Beyer, et Ruedi Aebersold. 2016. «On the Dependency of Cellular Protein Levels on mRNA Abundance». Cell 165 (3): 535-50. 10.1016/j.cell.2016.03.014.

Lu, Yi-Chien, Chad Sanada, Juliana Xavier-Ferrucio, Lin wang, Ping-Xia Zhang, H. Leighton Grimes, Meenakshi Venkatasubramanian, et al. 2018a. «The Molecular Signature of Megakaryocyte-Erythroid Progenitors Reveals a Role for the Cell Cycle in Fate Specification». Cell Reports 25 (8): 2083–2093.e4. 10.1016/j.celrep.2018.10.084.

Lu, Yi-Chien, Chad Sanada, Juliana Xavier-Ferrucio, Lin wang, Ping-Xia Zhang, H. Leighton Grimes, Meenakshi Venkatasubramanian, et al. 2018b. «The Molecular Signature of Megakaryocyte-Erythroid Progenitors Reveals a Role for the Cell Cycle in Fate Specification». Cell Reports 25 (8): 2083–2093.e4. 10.1016/j.celrep.2018.10.084.

Luecken, Malte D., M. Büttner, K. Chaichoompu, A. Danese, M. Interlandi, M. F. Mueller, D. C. Strobl, et al. 2022. «Benchmarking Atlas-Level Data Integration in Single-Cell Genomics». Nature Methods 19 (1): 41–50. 10.1038/s41592-021-01336-8.

Macosko, Evan Z., Anindita Basu, Rahul Satija, James Nemesh, Karthik Shekhar, Melissa Goldman, Itay Tirosh, et al. 2015. «Highly Parallel Genome-wide Expression Profiling of Individual Cells Using Nanoliter Droplets». Cell 161 (5): 1202–14. 10.1016/j.cell.2015.05.002.

Malumbres, Marcos. 2014. «Cyclin-Dependent Kinases». Genome Biology 15 (6): 122. 10.1186/gb4184.

Marino, Simeone, Ian B. Hogue, Christian J. Ray, et Denise E. Kirschner. 2008. «A Methodology for Performing Global Uncertainty and Sensitivity Analysis in Systems Biology». Journal of Theoretical Biology 254 (1): 178-96. 10.1016/j.jtbi.2008.04.011.

Masclef, Louis, Vanessa Dehennaut, Marlène Mortuaire, Céline Schulz, Maïté Leturcq, Tony Lefebvre, et Anne-Sophie Vercoutter-Edouart. 2019. «Cyclin D1 Stability Is Partly Controlled by O-GlcNAcylation». Frontiers in Endocrinology 10:106. 10.3389/fendo.2019.00106.

Mathieson, Toby, Holger Franken, Jan Kosinski, Nils Kurzawa, Nico Zinn, Gavain Sweetman, Daniel Poeckel, et al. 2018. «Systematic Analysis of Protein Turnover in Primary Cells». Nature Communications 9 (1): 689. 10.1038/s41467-018-03106-1.

Matsumoto, Akinobu, et Keiichi I. Nakayama. 2013. «Role of Key Regulators of the Cell Cycle in Maintenance of Hematopoietic Stem Cells». Biochimica et Biophysica Acta (BBA) - General Subjects 1830 (2): 2335-44. 10.1016/j.bbagen.2012.07.004.

Matsushime, H., M. F. Roussel, R. A. Ashmun, et C. J. Sherr. 1991. «Colony-Stimulating Factor 1 Regulates Novel Cyclins during the G1 Phase of the Cell Cycle». Cell 65 (4): 701–13. 10.1016/0092-8674(91)90101-4.

Mende, Nicole, Erika E. Kuchen, Mathias Lesche, Tatyana Grinenko, Konstantinos D. Kokkaliaris, Helmut Hanenberg, Dirk Lindemann, et al. 2015. «CCND1–CDK4–Mediated Cell Cycle Progression Provides a Competitive Advantage for Human Hematopoietic Stem Cells in Vivo». Journal of Experimental Medicine 212 (8): 1171–83. 10.1084/jem.20150308.

Miller, Donald M., Shelia D. Thomas, Ashraful Islam, David Muench, et Kara Sedoris. 2012. «C-Myc and Cancer Metabolism». Clinical Cancer Research: An Official Journal of the American Association for Cancer Research 18 (20): 5546-53. 10.1158/1078-0432.CCR-12-0977.

Muroyama, Yuki, et E. John wherry. 2021. «Memory T-Cell Heterogeneity and Terminology». Cold Spring Harbor Perspectives in Biology 13 (10): a037929. 10.1101/cshperspect.a037929.

Nakamura-Ishizu, Ayako, Hitoshi Takizawa, et Toshio Suda. 2014. «The Analysis, Roles and Regulation of Quiescence in Hematopoietic Stem Cells». Development 141 (24): 4656-66. 10.1242/dev.106575.

Ohtani, K., J. DeGregori, et J. R. Nevins. 1995. «Regulation of the Cyclin E Gene by Transcription Factor E2F1». Proceedings of the National Academy of Sciences of the United States of America 92 (26): 12146–50. 10.1073/pnas.92.26.12146.

Osorio, Daniel, et James J. Cai. 2021. «Systematic Determination of the Mitochondrial Proportion in Human and Mice Tissues for Single-Cell RNA-Sequencing Data Quality Control». Bioinformatics (Oxford, England) 37 (7): 963-67. 10.1093/bioinformatics/btaa751.

Ramezani-Rad, Parham, Cindi Chen, Zilu Zhu, et Robert C. Rickert. 2020. «Cyclin D3 Governs Clonal Expansion of Dark Zone Germinal Center B Cells». Cell Reports 33 (7): 108403. 10.1016/j.celrep.2020.108403.

Ranek, Jolene S., wayne Stallaert, J. Justin Milner, Margaret Redick, Samuel C. wolff, Adriana S. Beltran, Natalie Stanley, et Jeremy E. Purvis. 2024. «DELVE: Feature Selection for Preserving Biological Trajectories in Single-Cell Data». Nature Communications 15 (1): 2765. 10.1038/s41467-024-46773-z.

Riba, Andrea, Attila Oravecz, Matej Durik, Sara Jiménez, Violaine Alunni, Marie Cerciat, Matthieu Jung, Céline Keime, william M. Keyes, et Nacho Molina. 2022. «Cell Cycle Gene Regulation Dynamics Revealed by RNA Velocity and Deep-Learning». Nature Communications 13 (1): 2865. 10.1038/s41467-022-30545-8.

Rieckmann, Jan C., Roger Geiger, Daniel Hornburg, Tobias wolf, Ksenya Kveler, David Jarrossay, Federica Sallusto, et al. 2017. «Social Network Architecture of Human Immune Cells Unveiled by Quantitative Proteomics». Nature Immunology 18 (5): 583–93. 10.1038/ni.3693.

Robinson, Mark D., Davis J. McCarthy, et Gordon K. Smyth. 2010. «edgeR: A Bioconductor Package for Differential Expression Analysis of Digital Gene Expression Data». Bioinformatics (Oxford, England) 26 (1): 139-40. 10.1093/bioinformatics/btp616.

Robson, Samuel C., Lesley ward, Helen Brown, Heather Turner, Ewan Hunter, Stella Pelengaris, et Michael Khan. 2011. «Deciphering C-MYC-Regulated Genes in Two Distinct Tissues». BMC Genomics 12 (septembre):476. 10.1186/1471-2164-12-476.

Roussel, M. F., A. M. Theodoras, M. Pagano, et C. J. Sherr. 1995. «Rescue of Defective Mitogenic Signaling by D-Type Cyclins». Proceedings of the National Academy of Sciences of the United States of America 92 (15): 6837–41. 10.1073/pnas.92.15.6837.

Ruiz, Sergio, Athanasia D. Panopoulos, Aída Herrerías, Karl-Dimiter Bissig, Margaret Lutz, w. Travis Berggren, Inder M. Verma, et Juan Carlos Izpisua Belmonte. 2011. «A High Proliferation Rate Is Required for Cell Reprogramming and Maintenance of Human Embryonic Stem Cell Identity». Current Biology 21 (1): 45-52. 10.1016/j.cub.2010.11.049.

Sagot, Isabelle, et Damien Laporte. 2019. «The Cell Biology of Quiescent Yeast - a Diversity of Individual Scenarios». Journal of Cell Science 132 (1): jcs213025. 10.1242/jcs.213025.

Saleban, Mostafa, Erica L. Harris, et James A. Poulter. 2023. «D-Type Cyclins in Development and Disease». Genes 14 (7): 1445. 10.3390/genes14071445.

Sanidas, Ioannis, Robert Morris, Katerina A. Fella, Purva H. Rumde, Myriam Boukhali, Eric C. Tai, David T. Ting, Michael S. Lawrence, wilhelm Haas, et Nicholas J. Dyson. 2019. «A Code of Mono-Phosphorylation Modulates the Function of RB». Molecular Cell 73 (5): 985-1000.e6. 10.1016/j.molcel.2019.01.004.

Sawai, Catherine M., Jacquelyn Freund, Philmo Oh, Delphine Ndiaye-Lobry, Jamieson C. Bretz, Alexandros Strikoudis, Lali Genesca, et al. 2012. «Therapeutic Targeting of the Cyclin D3:CDK4/6 Complex in T Cell Leukemia». Cancer Cell 22 (4): 452–65. 10.1016/j.ccr.2012.09.016.

Sewing, A., C. Bürger, S. Brüsselbach, C. Schalk, F. C. Lucibello, et R. Müller. 1993. «Human Cyclin D1 Encodes a Labile Nuclear Protein whose Synthesis Is Directly Induced by Growth Factors and Suppressed by Cyclic AMP». Journal of Cell Science 104 ( Pt 2) (février):545-55. 10.1242/jcs.104.2.545.

Sherr, C. J. 1996. «Cancer Cell Cycles». Science (New York, N.Y.) 274 (5293): 1672–77. 10.1126/science.274.5293.1672.

Sherr, C. J., et J. M. Roberts. 1999. «CDK Inhibitors: Positive and Negative Regulators of G1-Phase Progression». Genes & Development 13 (12): 1501–12. 10.1101/gad.13.12.1501.

Shreeram, Sathyavageeswaran, weng Kee Hee, et Dmitry V. Bulavin. 2008. «Cdc25A Serine 123 Phosphorylation Couples Centrosome Duplication with DNA Replication and Regulates Tumorigenesis». Molecular and Cellular Biology 28 (24): 7442-50. 10.1128/MCB.00138-08.

Signer, Robert A. J., Jeffrey A. Magee, Adrian Salic, et Sean J. Morrison. 2014. «Haematopoietic Stem Cells Require a Highly Regulated Protein Synthesis Rate». Nature 509 (7498): 49-54. 10.1038/nature13035.

Skotheim, Jan M., Stefano Di Talia, Eric D. Siggia, et Frederick R. Cross. 2008. «Positive Feedback of G1 Cyclins Ensures Coherent Cell Cycle Entry». Nature 454 (7202): 291-96. 10.1038/nature07118.

Spencer, Sabrina L., Steven D. Cappell, Feng-Chiao Tsai, K. wesley Overton, Clifford L. wang, et Tobias Meyer. 2013. «The Proliferation-Quiescence Decision Is Controlled by a Bifurcation in CDK2 Activity at Mitotic Exit». Cell 155 (2): 369-83. 10.1016/j.cell.2013.08.062.

Sukys, Augustinas, et Ramon Grima. 2025. «Cell-Cycle Dependence of Bursty Gene Expression: Insights from Fitting Mechanistic Models to Single-Cell RNA-Seq Data». Nucleic Acids Research 53 (7): gkaf295. 10.1093/nar/gkaf295.

Sun, Xiaoming, Aizhan Bizhanova, Timothy D. Matheson, Jun Yu, Lihua Julie Zhu, et Paul D. Kaufman. 2017. «Ki-67 Contributes to Normal Cell Cycle Progression and Inactive X Heterochromatin in P21 Checkpoint-Proficient Human Cells». Molecular and Cellular Biology 37 (17): e00569-16. 10.1128/MCB.00569-16.

Surmacz, E., K. Reiss, C. Sell, et R. Baserga. 1992. «Cyclin D1 Messenger RNA Is Inducible by Platelet-Derived Growth Factor in Cultured Fibroblasts». Cancer Research 52 (16): 4522–25.

Swaffer, Matthew P., Jacob Kim, Devon Chandler-Brown, Maurice Langhinrichs, Georgi K. Marinov, william J. Greenleaf, Anshul Kundaje, Kurt M. Schmoller, et Jan M. Skotheim. 2021. «Transcriptional and Chromatin-Based Partitioning Mechanisms Uncouple Protein Scaling from Cell Size». Molecular Cell 81 (23): 4861-4875.e7. 10.1016/j.molcel.2021.10.007.

Teierle, Samantha M., Ying Huang, Adam S. Kittai, Seema A. Bhat, Michael Grever, Kerry A. Rogers, weiqiang Zhao, et al. 2023. «Characteristics and Outcomes of Patients with CLL and CDKN2A/B Deletion by Fluorescence in Situ Hybridization». Blood Advances 7 (23): 7239–42. 10.1182/bloodadvances.2023010753.

«The Chemical Basis of Morphogenesis». 1952. Philosophical Transactions of the Royal Society of London. Series B, Biological Sciences 237 (641): 37–72. 10.1098/rstb.1952.0012.

Thomas, Lance R., et william P. Tansey. 2011. «Proteolytic Control of the Oncoprotein Transcription Factor Myc». Advances in Cancer Research 110:77-106. 10.1016/B978-0-12-386469-7.00004-9.

Vercruysse, Jasmien, Alexandra Baekelandt, Nathalie Gonzalez, et Dirk Inzé. 2020. «Molecular Networks Regulating Cell Division during Arabidopsis Leaf Growth». Journal of Experimental Botany 71 (8): 2365-78. 10.1093/jxb/erz522.

Vigo, E., H. Müller, E. Prosperini, G. Hateboer, P. Cartwright, M. C. Moroni, et K. Helin. 1999. «CDC25A Phosphatase Is a Target of E2F and Is Required for Efficient E2F-Induced S Phase». Molecular and Cellular Biology 19 (9): 6379–95. 10.1128/MCB.19.9.6379.

wang, Zhaoming, et Jinghui Zhang. 2025. «Genetic and Epigenetic Bases of Long-Term Adverse Effects of Childhood Cancer Therapy». Nature Reviews Cancer 25 (2): 129-44. 10.1038/s41568-024-00768-6.

wei, Taiyun, et Viliam Simko. 2024. R package «corrplot»: Visualization of a Correlation Matrix. https://github.com/taiyun/corrplot.

weinberg, R. A. 1995. «The Retinoblastoma Protein and Cell Cycle Control». Cell 81 (3): 323–30. 10.1016/0092-8674(95)90385-2.

wiśniewski, Jacek R., Marco Y. Hein, Jürgen Cox, et Matthias Mann. 2014. «A “Proteomic Ruler” for Protein Copy Number and Concentration Estimation without Spike-in Standards». Molecular & Cellular Proteomics: MCP 13 (12): 3497-3506. 10.1074/mcp.M113.037309.

wolf, F. Alexander, Philipp Angerer, et Fabian J. Theis. 2018. «SCANPY: Large-Scale Single-Cell Gene Expression Data Analysis». Genome Biology 19 (1): 15. 10.1186/s13059-017-1382-0.

wu, Lizhao, Cynthia Timmers, Baidehi Maiti, Harold I. Saavedra, Ling Sang, Gabriel T. Chong, Faison Nuckolls, et al. 2001. «The E2F1–3 Transcription Factors Are Essential for Cellular Proliferation». Nature 414 (6862): 457–62. 10.1038/35106593.

Xia, Lucy, Christy Lee, et Jingyi Jessica Li. 2024. «Statistical Method scDEED for Detecting Dubious 2D Single-Cell Embeddings and Optimizing t-SNE and UMAP Hyperparameters». Nature Communications 15 (1): 1753. 10.1038/s41467-024-45891-y.

Zatulovskiy, Evgeny, Shuyuan Zhang, Daniel F. Berenson, Benjamin R. Topacio, et Jan M. Skotheim. 2020. «Cell Growth Dilutes the Cell Cycle Inhibitor Rb to Trigger Cell Division». Science (New York, N.Y.) 369 (6502): 466-71. 10.1126/science.aaz6213.

Zhang, Shuyuan, Lucas Fuentes Valenzuela, Evgeny Zatulovskiy, Lise Mangiante, Christina Curtis, et Jan M. Skotheim. 2024. «The G1/S Transition Is Promoted by Rb Degradation via the E3 Ligase UBR5». bioRxiv: The Preprint Server for Biology, avril, 2023.10.03.560768. 10.1101/2023.10.03.560768.

Zhang, Xuan, Baobao Song, Maximillian J. Carlino, Guangyuan Li, Kyle Ferchen, Mi Chen, Evrett N. Thompson, et al. 2024. «An Immunophenotype-Coupled Transcriptomic Atlas of Human Hematopoietic Progenitors». Nature Immunology 25 (4): 703–15. 10.1038/s41590-024-01782-4.

Zhou, Yu, Yan Geng, Yujiao Zhang, Yubin Zhou, Chen Chu, Samanta Sharma, Anne Fassl, Deborah Butter, et Piotr Sicinski. 2020. «The Requirement for Cyclin E in C-Myc Overexpressing Breast Cancers». Cell Cycle (Georgetown, Tex.) 19 (20): 2589-99. 10.1080/15384101.2020.1804720.

Zikry, Tarek M., Samuel C. wolff, Jolene S. Ranek, Harris M. Davis, Ander Naugle, Namit Luthra, Austin A. whitman, et al. 2024. «Cell Cycle Plasticity Underlies Fractional Resistance to Palbociclib in ER+/HER2-Breast Tumor Cells». Proceedings of the National Academy of Sciences of the United States of America 121 (7): e2309261121. 10.1073/pnas.2309261121.

